# Identification of subfunctionalized aggregate-remodeling J-domain proteins in plants

**DOI:** 10.1101/2020.10.14.340331

**Authors:** Yogesh Tak, Silviya S. Lal, Shilpa Gopan, Madhumitha Balakrishnan, Amit K. Verma, Sierra J. Cole, Rebecca E. Brown, Rachel E. Hayward, Justin K. Hines, Chandan Sahi

## Abstract

Hsp70s and J-domain proteins (JDPs) are among the most critical components of the cellular protein quality control machinery, playing crucial roles in preventing and solubilizing cytotoxic protein aggregates. Bacteria, yeast and plants additionally have large, multimeric Hsp100-class disaggregases which, allow the resolubilization of otherwise “dead-end” aggregates, including amyloids. JDPs interact with aggregated proteins and specify the aggregate remodeling activities of Hsp70s and Hsp100s. Plants have a complex network of cytosolic Hsp70s and JDPs, however the aggregate remodeling properties of plant JDPs are not well understood. Here we identify evolutionary-conserved Class II JDPs in the model plant *Arabidopsis thaliana* with distinct aggregate remodeling functionalities. We identify eight plant orthologs of the yeast protein, Sis1, the principal JDP responsible for directing the yeast chaperone machinery for remodeling protein aggregates. Expression patterns vary dramatically among the eight paralogous proteins under a variety of stress conditions, indicating their subfunctionalization to address distinct stressors. Consistent with a role in solubilizing cytotoxic protein aggregates, six of these plant JDPs associate with heat-induced protein aggregates *in vivo* as well as colocalize with plant Hsp101 to distinct heat-induced protein aggregate centers. Finally, we show that these six JDPs can differentially remodel multiple model protein aggregates in yeast confirming their involvement in aggregate resolubilization. These results demonstrate that compared to complex metazoans, plants have a robust network of JDPs involved in aggregate remodeling activities with the capacity to process a variety of protein aggregate conformers.

## Introduction

All organisms face the challenge of maintaining cellular proteostasis for proper growth and development in the context of constant and diverse stressors. Proteostasis ensures the balance between numerous cellular processes, including protein synthesis, folding, transport, aggregation, and turnover, thereby maintaining the overall fidelity of cellular proteome (1). Proteins can misfold for a plethora of reasons, including cellular stresses, mutations, defects in protein synthesis or sorting, or chaperone overload (2). Misfolded proteins can form abnormal contacts with other proteins and seed the formation of larger protein aggregates, which are often cytotoxic (3). Broadly, these aggregates are either amorphous deposits or ordered amyloid fibrils (4, 5). Either of these aggregates can further disrupt cellular proteostasis, which can hamper vital cellular processes (6).

Being sessile, plants have an increased burden to continuously integrate their cellular physiology with diverse environmental cues to maintain proper growth and development as well as safeguard reproductive success. Protein homeostasis in plants is vulnerable to adverse biotic or abiotic stress conditions that are detrimental to plant health and can ultimately lead to a decrease in productivity (7). High-temperature stress is among the primary factors causing proteotoxic stress, eventually leading to adverse physiological consequences in plants (8, 9). To counter stress-induced challenges, plants are equipped with a vast network of heat-shock proteins (Hsps), often referred to as molecular chaperones (10). Molecular chaperones assist in protein folding, prevention of misfolding and aggregation, reactivation of aggregates, and degradation or sequestration of problematic protein substrates (5, 11, 12).

Small heat shock proteins (sHsps), Hsp40s, Hsp70s, and Hsp100s (when present) play vital roles in liberating and refolding trapped polypeptides from protein aggregates (5). sHsps work as the first line of defense against proteotoxic stress, as they bind to misfolded proteins in an ATP-independent fashion and prevent irreversible aggregation (13). Moreover, denatured protein substrates bound to sHsps remain in a folding competent state so that Hsp70s, with their obligate cochaperone Hsp40s (alternatively and hereafter called J-domain proteins or JDPs) can reactivate them (14). Additionally, in bacteria, yeast and plants the Hsp70: JDP system further cooperates with AAA+ ATPases of the Caseinolytic Protease B/heat shock protein 100 (ClpB/Hsp100) family in a synergic, ATP-driven, bi-chaperone system for disaggregation, in which the JDP acts as a cochaperone and deliver substrates to Hsp70 and therefore Hsp100 (15–17). The disaggregation activities of Hsp100s have been conserved with similar proteins found in bacteria (ClpB), yeast (Hsp104), and plants (Hsp101) (18–20). sHsps, JDPs, Hsp70s, and Hsp101 have been shown to have a significant role in stress sensing and tolerance (21–26). Studies have implicated sHsps and Hsp101 in aggregate remodeling in plants (21, 27), however there is no such report on plant JDPs.

JDPs are crucial co-chaperones that define the substrate specificities of Hsp70 and Hsp100 (17, 28, 29). JDPs are a highly heterogeneous group of proteins. All JDPs have a conserved J-domain, which is critical for the stimulation of the otherwise weak ATPase activity of their partner Hsp70s (15). Based on their domain organization and similarity to DnaJ of *E. coli*, JDPs have been divided into three classes, Class I, II and III (30). Among these, the Class II JDPs, including Sis1 in yeast, DNAJB1 in mammals and DROJ1 in flies have been implicated in aggregate remodelling (31–33). The aggregate remodelling activities of Sis1 are best understood. Sis1 is an essential cytosolic Class II JDP that contains a J-domain, a glycine-rich central region (G/F-G/M), a carboxyl-terminal region (CTD), which has a client binding pocket, and dimerization domain (DD) (34). Sis1 is required for the remodeling of different protein aggregates including terminally misfolded proteins and prion (31, 35–38). Prions are self-replicating proteinaceous infectious particles. One characteristic feature of all prions is that they have the ability to convert the cellular pool of the prion protein into a misfolded form, resulting into aggregation (39). Among all 22 JDPs in yeast, Sis1 uniquely cooperates with Hsp70 and Hsp104 in fragmenting prion aggregates such as [*PSI*^+^] and [*RNQ*^+^] to generate seeds, an essential step in the maintenance of prions in yeast cells (17, 31, 40). The [*PSI*^+^] prion is formed of the Sup35 protein, a translation termination factor. In [*PSI*^+^] cells, Sup35 is sequestered in amyloid aggregates, leading to the nonsense-suppression phenotype. The [*PSI*^+^] prion has different heritable states which result from amyloid polymorphisms (alternative conformers) (41). These are termed “prion strains” in mammalian systems or “variants” in yeast. [*PSI*^+^] variants are classified as “weak” and “strong” based on the strength of the nonsense suppression phenotype, which results from differences in the soluble pool of Sup35. Variants with distinct aggregate sizes and number of prion seeds per cell lead to differential amounts of unincorporated Sup35 which alters translational read through efficiencies (42). In yeast strains with a nonsense mutation in the adenosine synthesis pathway, [*psi*^-^] strains form red colonies due to the accumulation of a red-pigmented intermediate. Cells bearing strong [*PSI*^+^] variants, in contrast, grow into white or light pink colonies, while cells with weak [*PSI*^+^] variants form dark pink colonies due to translational readthrough and partial adenosine synthesis. Apart from [*PSI*^+^] maintenance, Sis1 is the only factor other than Hsp70 and Hsp104 that is known to be required for the maintenance of the yeast prion [*RNQ*^+^] (35, 38). Previous reports suggest that the human (Hdj1), and Drosophila (Droj1) ortholog of Sis1 can maintain [*RNQ*^+^] and strong [*PSI*^+^] prions in budding yeast (31, 32, 43).

To identify and functionally characterize aggregate remodeling JDPs in plants, we chose the orthologs of Sis1 in the model plant *Arabidopsis thaliana*. Previously we showed that the *A. thaliana* Class II JDP, atDjB1 could substitute for Sis1in budding yeast (44). Besides rescuing the lethality of *sis1Δ* strain, atDjB1 could also maintain the commonly-studied yeast prions, [*RNQ*^+^] and [*PSI*^+^]. Interestingly, Arabidopsis has at least seven more Sis1-like JDPs (atDjBs). However, the aggregate remodeling properties of these JDPs have not been characterized. Here we show that atDjBs can associate with heat-induced protein aggregates and co-localize with Hsp101 disaggregase, the marker for heat-denatured protein aggregates in isolated protoplasts. In the absence of an assayable loss of function phenotype in Arabidopsis and lack of model protein aggregate substrates in plants we characterized the functional specificities and aggregate-remodeling properties of six of these plant JDPs using yeast genetic tools. Selected, well characterized yeast prions were exploited as model protein aggregates, as these have been immensely useful in understanding the aggregate remodeling functions of chaperones including JDPs (45, 46). The six tested atDjBs not only complement essential functions of Sis1 but also differentially remodel yeast prions. Put together, our results indicate that plants have an elaborate network of aggregate remodeling chaperones. Moreover, the aggregate remodeling properties of JDPs seem to be subfunctionalized to provide better adaptation to land plants during diverse environmental conditions.

## Results

### atDjBs are highly similar to Sis1

We previously found eight orthologs of Sis1 in *A. thaliana*, namely, atDjB1, atDjB2, atDjB3, atDjB4, atDjB5, atDjB6, atDjB10, and atDjB17 (44). These atDjBs not only share the domain organization of Sis1 but also exhibit high sequence similarity among one another (68.1%–89.4%), with Sis1 (49.20%–53.9%), and with the human ortholog, Hdj1 (57.5%–63.7%). (Fig. 1A; 1B; S1; Table 1). The e-values range between 1.00E-61 (atDjB1) to 2.00E-52 (atDjB17) obtained by NCBI BLASTp analyses using *S. cerevisiae* Sis1 as a query sequence (Fig. S2A). Further, the C-terminal domains of all atDjBs protein share high degree of similarity to Sis1 (Fig. S2B). Notably, even the hydrophobic client binding pocket is highly conserved among all eight JDPs (Fig. S2C).

**Fig. 1.**
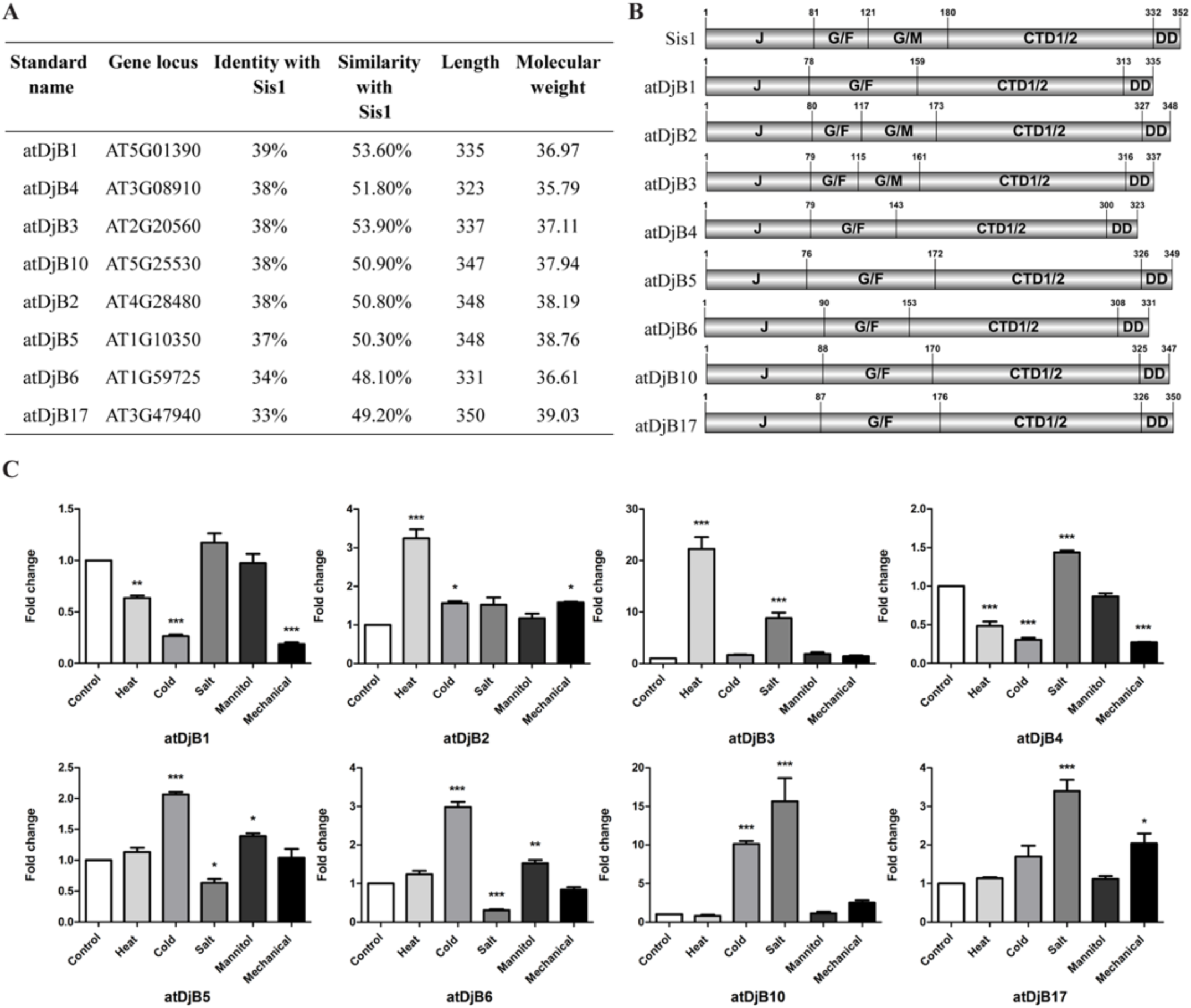
Protein sequence features of *A. thaliana* atDjBs and their expression profiling. **A.** Percent identity and similarity between full-length protein sequences of *A. thaliana* atDjBs and Sis1. **B.** Domain organization of *A. thaliana* atDjBs. J-domain (J), glycine phenylalanine-rich region (GF), glycine methionine-rich region (GM), C-terminal peptide-binding domains I and II (CTD1/2), dimerization domain (DD). The domain structures were drawn to scale, and amino-acid numbers were shown over each domain. **C.** Differential expression pattern of atDjBs under abiotic stress conditions. qRT-PCR analysis showing relative expression of atDjBs examined in the WT under control and different abiotic stress conditions: Heat (37°C for 60min), Cold (4°C for 24h), Salt (150mM NaCl for 24h), Mannitol (150mM Mannitol for 24h), or Mechanical (mechanical injury, 60min). Gene expression levels were normalized in stressed samples with the respective genes in unstressed samples, followed by normalization with the expression level of *ACTIN* (AT3G1870) gene. Significant transcript level differences were indicated by asterisks (*** *P<0.0001*, ** *P<0.001*, * *P<0.01*) analyzed using 2way-ANOVA with Dunnett’s multiple comparison test. Data are mean ±SD of three biological and technical replicates.

**Table 1.**
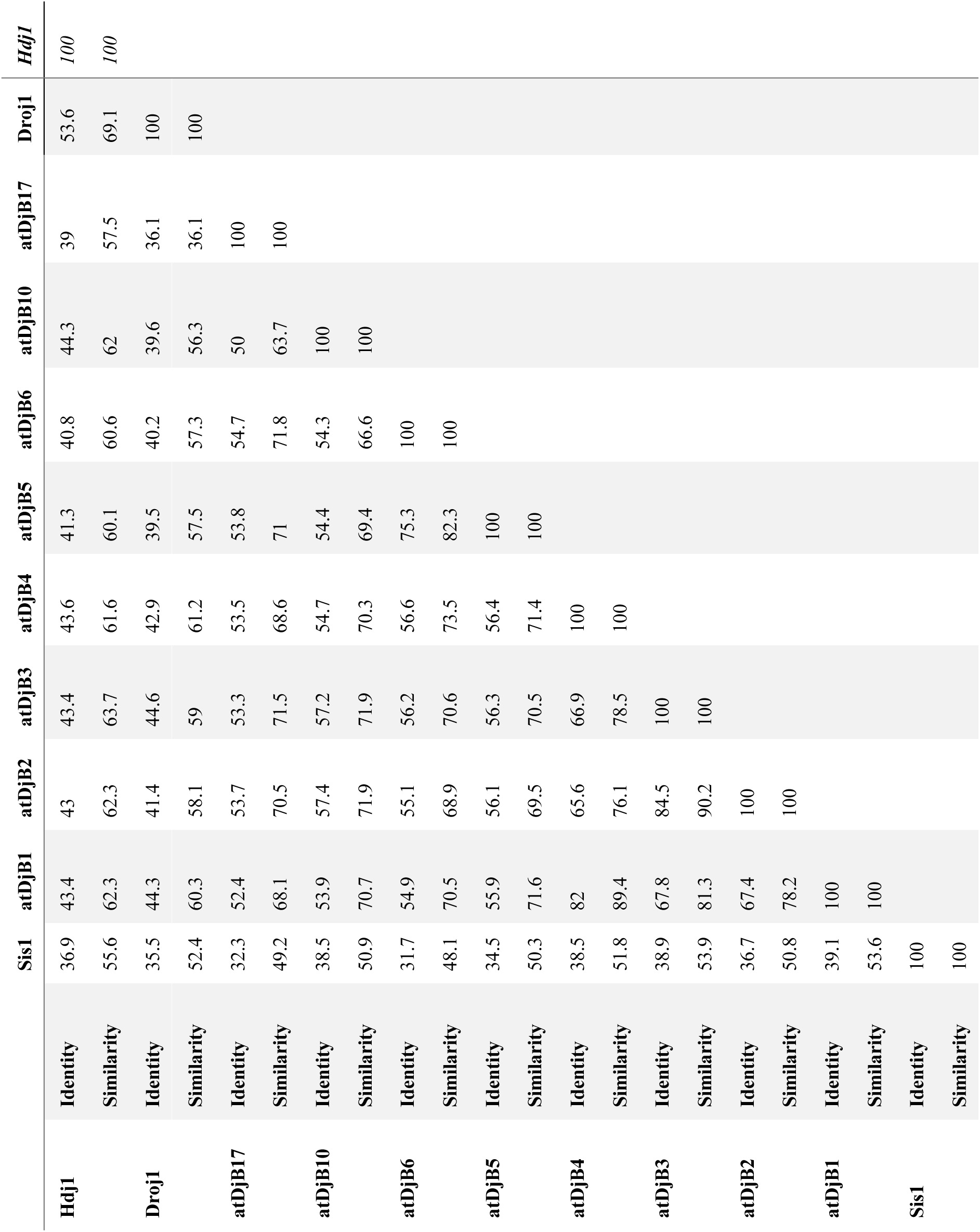
Percent amino acid sequence similarity and identity among *A. thaliana*atDjBs, Sis1, Droj1 and Hdj1. Full length protein sequences were used to calculate the similarity and identity with EMBOSS Needle online tool.

### *atDjBs* are expressed differentially

To understand the functional significance of these atDjBs in *A. thaliana*, we first analyzed their expression under different abiotic stress conditions through quantitative RT-PCR (Fig. 1C) (44). Interestingly, only atDjB2 and atDjB3 were significantly upregulated under heat stress (37°C for 60 min), atDjB3, atDjB4, atDjB10, and atDjB17 were induced by salt stress (150 mM NaCl for 24 h), while atDjB5, atDjB6, and atDjB10 were cold stress (4°C for 24 h) regulated. Similarly, atDjB1 and atDjB4 showed significant down-regulation under heat, cold, and salt stress, and atDjB6 only under salt stress. Our results show that the atDjBs are expressed differentially and even though highly similar, these proteins might have different stress associated functionalities in *A. thaliana*.

### atDjBs interact with misfolded or aggregated proteins

Under high temperature stress, a large portion of the plant proteome faces the risk of misfolding and aggregation (47). To test whether atDjBs are involved in heat stress response, we looked at the atDjB1 protein levels in total protein fraction of unstressed, acclimatized, and heat-stressed samples. A band corresponding to atDjB1 was detected in unstressed (23°C) conditions, which was upregulated upon acclimatization (38°C for 90 min, followed by 120 min at 23°C) and under heat stress (180 min at 45°C with preconditioning) (Fig. 2). However, the polyclonal antibody used in this assay was raised against atDjB1-CTD and cross-reacts with atDjB2, atDjB3, and atDjB4 (Fig. S3), so at this time we cannot rule out the possibility that the band visible on the western blots represents the heat-inducible orthologs, atDjB2 and atDjB3 or other atDjBs as well.

**Fig. 2.**
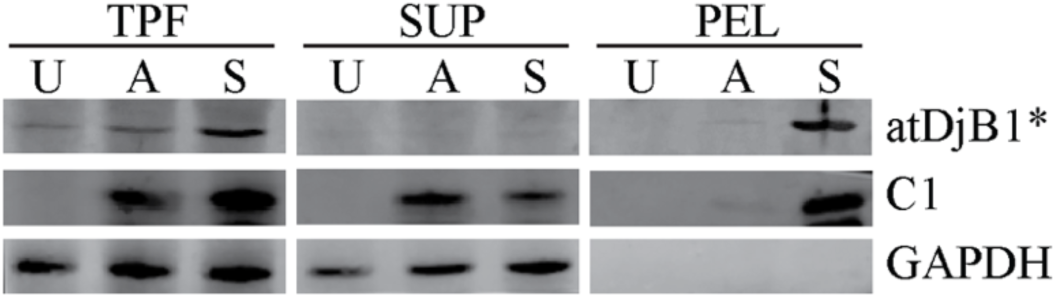
atDjBs are heat-inducible and associate with heat-induced protein aggregates. Total protein lysate was isolated from Arabidopsis seedlings that were unstressed (U), acclimatized at 38°C for 90 min followed by 120 min at 22°C (A), and heat stressed for 180min at 45°C with preconditioning (S). Lysates were then fractionated by centrifugation into supernatant (SUP) and pellet (PEL) fractions, and resolved along with total protein fraction (TPF) on 15% SDS-PAGE and electroblotted. Blots were probed with anti-atDjB1, anti-sHsp antibodies and anti-GAPDH as a loading control. Asterisk (*) represent the polyclonal antibody against atDjB1 that cross reacts with atDjB2, atDjB3 and atDjB4.

Since some JDPs are known to prevent irreversible aggregation of heat-denatured proteins by interacting with them, we asked if atDjBs have a similar role in remodeling heat-denatured proteins in *A. thaliana*. To test this, we first sought to identify the association of atDjBs with the heat-denatured proteins *in planta*. We assessed the presence of atDjBs in either soluble or pellet fractions under control conditions or after acclimatization and heat stress. During heat stress, atDjB(s) partitioned into the insoluble fraction similar to the sHSP C1, a known marker for heat-induced protein aggregates (Fig. 2). As expected, GAPDH was detected only in the soluble fractions of all samples, which perfectly aligns with a previous report (21). This result suggests that atDjBs interact with the heat-denatured protein aggregates in cells.

### atDjBs colocalize with Hsp101 to distinct heat-induced punctae in *A. thaliana* protoplasts

To individually test the ability of atDjBs to associate with heat-denatured protein aggregates, we generated C-terminal GFP-tagged constructs of atDjB1–6. Even after repeated efforts, we could not clone the full-length cDNAs of atDjB10 and atDjB17, so these were not included in any further studies. atDjB1–6-GFP fusion proteins were transiently expressed in *A. thaliana* protoplasts, and their subcellular localization was assessed using confocal laser scanning microscopy at 23°C, or after heat stress for either 30 min at 38°C or 15 min at 45°C. Under control conditions, atDjB1–6 exhibited a diffuse fluorescence pattern (Fig. 3). In contrast, heat stress resulted in a transition from diffuse to punctate cellular distribution (Fig. S4A; S4B).

**Fig. 3.**
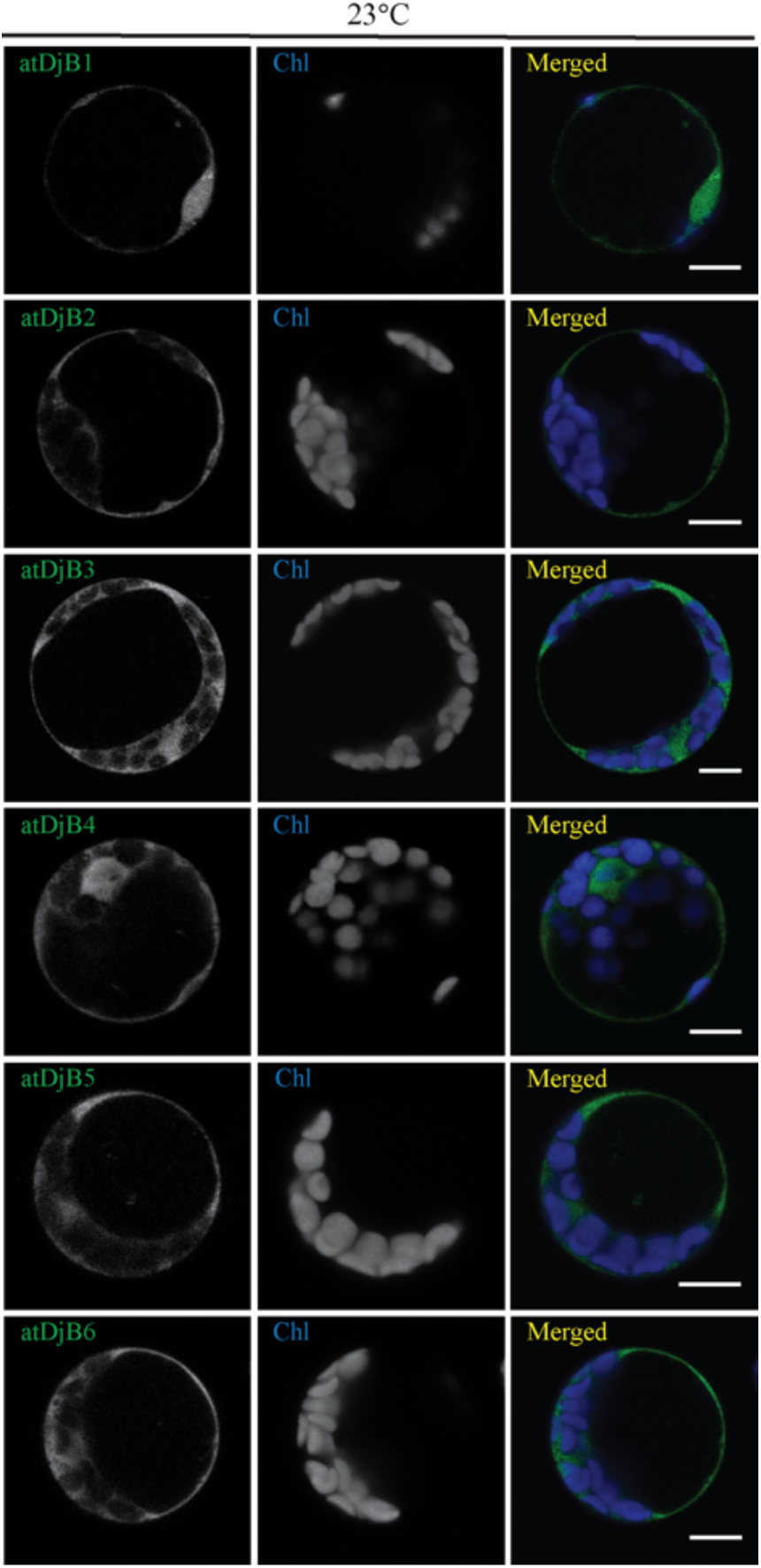
Subcellular localization of atDjBs in *A. thaliana* protoplasts. *A. thaliana* leaf protoplasts were transfected with the vectors expressingatDjB1-GFP, atDjB2-GFP, atDjB3-GFP, atDjB4-GFP, atDjB5-GFP, and atDjB6-GFP fusion proteins under control conditions (23°C). The green signals indicate GFP, and the blue signals indicate autofluorescence of chlorophyll. GFP-mediated fluorescence, derived from individual atDjBs and chlorophyll autofluorescence were visualized using FITC (495nm/519nm) and Cy5 (678nm/694nm) filter set. All images are single slice of confocal sections taken with living protoplast 12 h after transfection. Scale bar, 10 μm.

Hsp101 is the major disaggregase of the Hsp100 class in *A. thaliana* that is known to solubilize heat-induced aggregates and improve thermotolerance (21). Upon heat stress, Hsp101 forms multiple cytosolic puncta and, thus, has been used as a marker for protein aggregate centers (PACs)in plant cells (27, 48). Hsp104, the yeast ortholog of Arabidopsis Hsp101, works with the Hsp70: JDP chaperone machinery to remodel multiple types of aggregates (17, 49). Previously, we showed that atDjB1 could remodel yeast model aggregate proteins in place of Sis1, demonstrating its ability to cooperate with yeast Hsp70 in aggregate remodeling (44). Because atDjBs fractionated with heat-denatured aggregates in our centrifugation assays, we hypothesized that atDjBs studied here likely to have a role in aggregate remodeling in plants, and if so, they should colocalize with Hsp101 at PACs during heat stress. To test this, we generated an N-terminal RFP-tagged Hsp101 construct and co-transformed *A. thaliana* protoplasts with plasmids expressing individual atDjB-GFP constructs. We focused only on heat stress treatment at 38°C for 30min to check the behavior of the proteins, as 15 min at 45°C resulted in several poly-disperse punctae that were not easily differentiable from each other (Fig. S5B). Similarly, during the control (23°C) condition, fluorescence was not differentiable as all proteins analyzed displayed a diffuse cytosolic distribution in protoplasts (Fig. S5A). As shown by the merged confocal fluorescence images and statistical analysis, atDjB1–4 and atDjB6 colocalized with Hsp101 at distinct punctate structures in Arabidopsis protoplasts following heat stress at 38°C for 30 min (Fig. 4A). Unlike other atDjBs, atDjB5 didn’t show significant co-localization with Hsp101-RFP, although it also accumulated at distinct punctate structures (Fig. 4B). These results suggest that, upon high-temperature, atDjBs are distributed to distinct heat-induced aggregates along with Hsp101 at PACs, indicating that they likely work with Hsp101 to remodel and/or solubilize heat-denatured protein aggregates in *A. thaliana*.

**Fig. 4.**
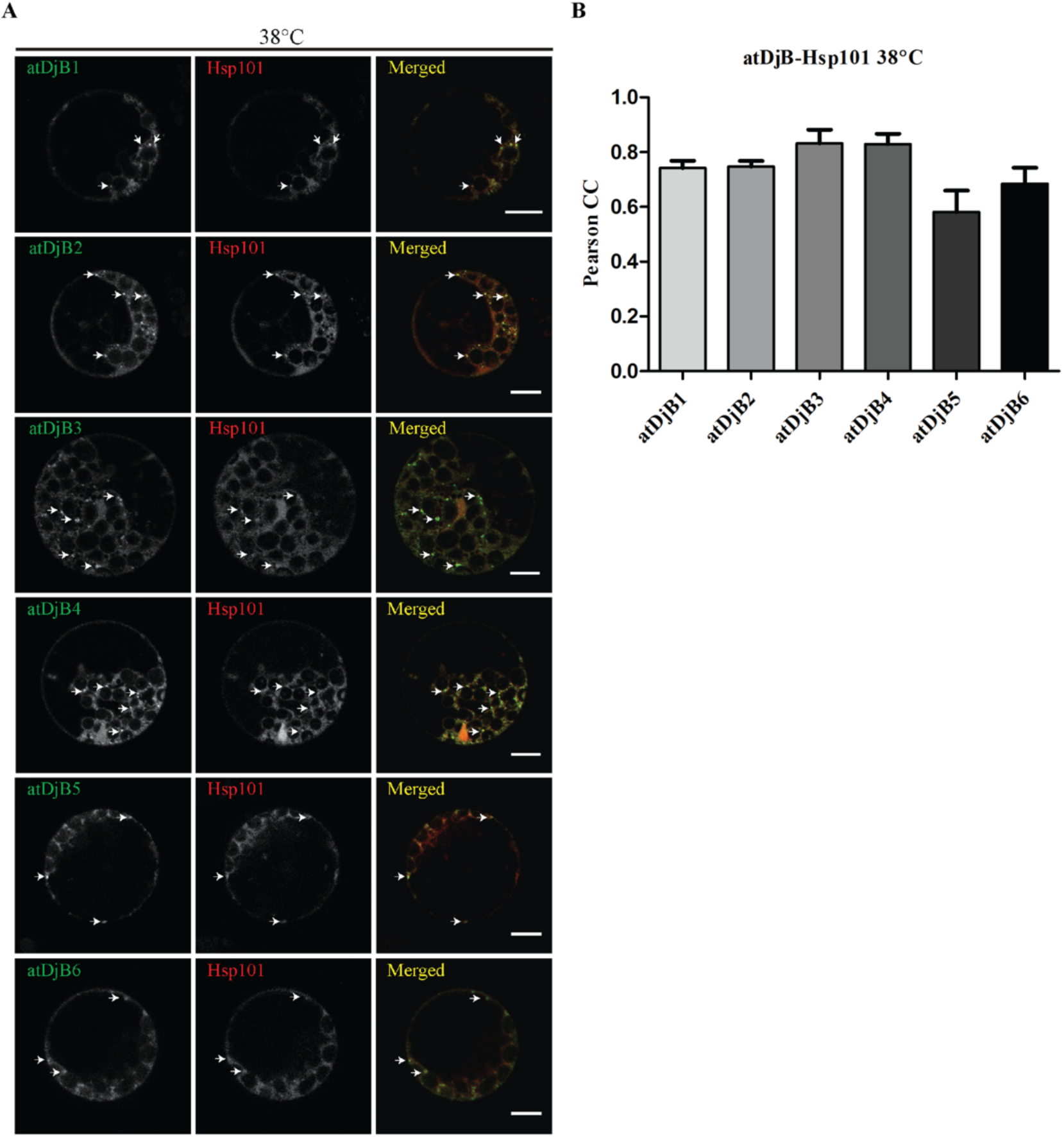
Heat stress induced multiple cytoplasmic foci of atDjBs colocalize to Hsp101 foci in *A. thaliana* protoplast. **A.** Protoplast of *A. thaliana* transiently expressing individual atDjB-GFP constructs and Hsp101-RFP were preconditioned at 38°C for 30 min. Green indicates GFP, red indicates RFP, and blue indicates the autofluorescence of chlorophyll. GFP-mediated fluorescence, derived from individual atDjBs, RFP-mediated fluorescence derived from Hsp101, and chlorophyll autofluorescence were visualized using FITC (495nm/519nm), m-RFP (572nm/606nm) and Cy5 (678nm/694nm) filter set. All images are single slice of confocal sections taken with living protoplast immediately after the above mentioned stress treatment. Scale bar, 10 μm. **B.** Quantification of colocalization of atDjBs (GFP signal) with Hsp101 (RFP-signal) by Pearson’s correlation coefficient, each bar represents the average correlation coefficient of (±SD) minimum five different protoplasts. Arrow (→) represent the cytoplasmic foci for the individual atDjB’s.

### atDjB1–6 can substitute for Sis1 in budding yeast

To further explore the functional similarity and diversity among the six atDjB paralogs, we took advantage of the unicellular eukaryote *Saccharomyces cerevisiae*, in which, as noted above, the role of Sis1 in protein aggregate remodeling is well characterized. We first tested if atDjB1–6 rescue the viability of a *sis1Δ* strain. For this, full-length cDNAs of atDjB1–6 were cloned in vectors with a constitutive *TEF1* promoter (50). A *sis1Δ* yeast strain harboring*SIS1* from a *URA3* plasmid (*URA3*-*SIS1*) was transformed with plasmids expressing individual atDjB1–6 proteins. Transformants were grown on medium containing 5-FOA to counterselect against the *URA3-SIS1* plasmid, and thus maintain only the plasmid expressing individual atDjB proteins. Results showed that like atDjB1, other five atDjBs could also substitute for Sis1 in yeast (Fig. 5A). Importantly, this result also implies that all six plant proteins can regulate the ATPase activity of yeast Hsp70s. Next, we assessed the growth of *sis1Δ* cells expressing different atDjBs at different temperatures. As shown in (Fig. 5B), atDjB1-4 rescued growth of *sis1Δ* cells similarly to Sis1. However, atDjB5 and atDjB6 could not support growth at 34°C (Fig. 5B). This partial complementation by atDjB5 and atDjB6 was most likely due to functional differences rather than protein expression, as evidenced by comparable expression levels of all the tested 1X-HA tagged atDjB proteins (Fig. 5C).

**Fig. 5.**
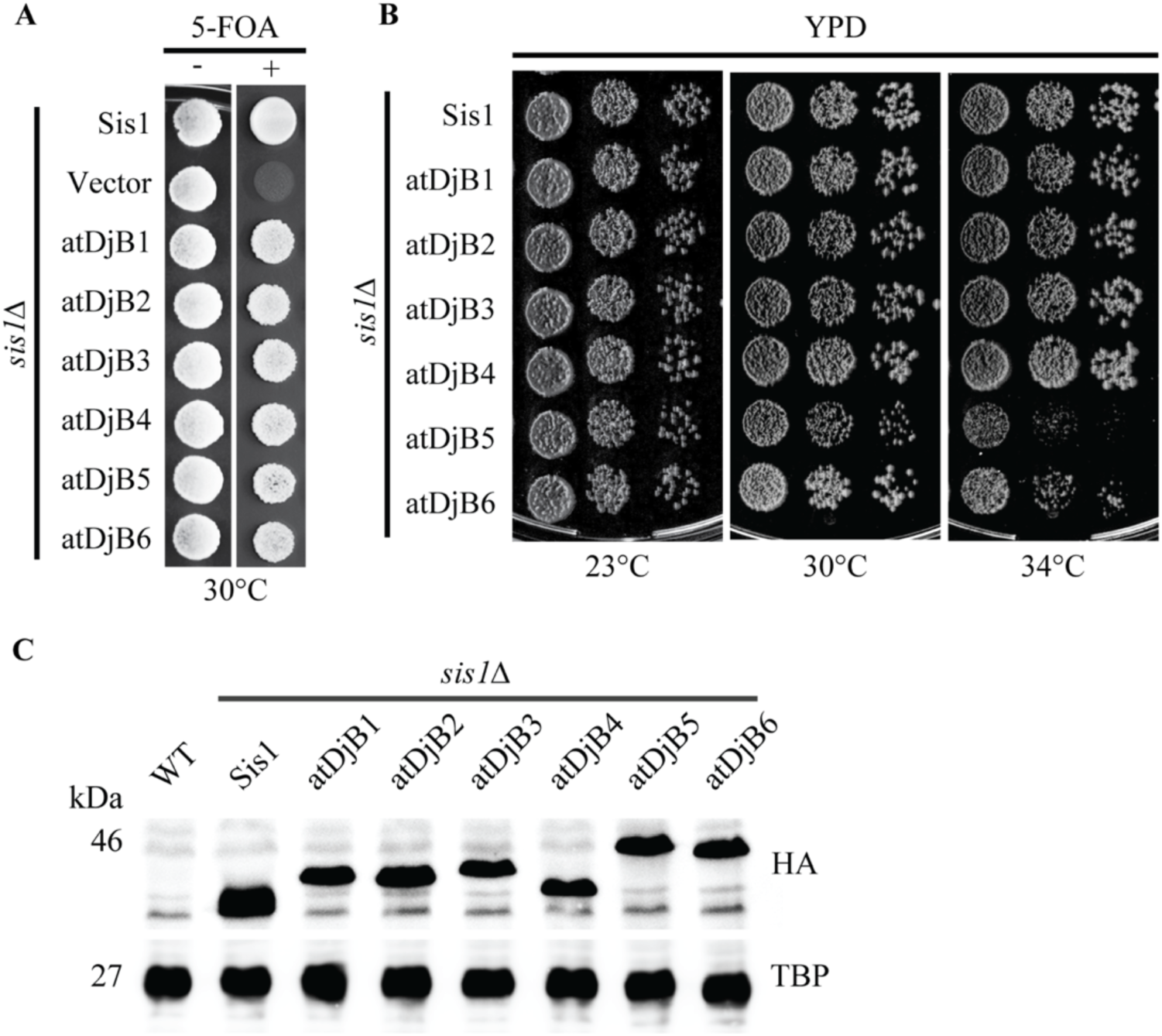
atDjB1-6 rescue the essential functions of Sis1 in *S. cerevisiae*. **A.** Equal volume of ten-fold serial dilutions of *sis1Δ* [*URA3*-*SIS1*] cells harboring empty pRS413 plasmid (vector) or pRS413 expressing Sis1 or atDjB1-6 were spotted on media with (+) or without (-) 5-fluoroorotic acid (5-FOA) and incubated at 30°C for 3 days. **B.** Equal volume of ten-fold serial dilutions of *sis1Δ* cells harboring pRS413-Sis1 (Sis1) or pRS413 expressing atDjB1-6 were spotted on YPD plates and incubated at indicated temperatures for 3 days. **C.** Equal amounts of total cell lysate prepared from s*is1Δ* cells harboring plasmids expressing HA-tagged constructs of Sis1 or atDjB1-6 were resolved on SDS-PAGE, electroblotted, and probed with anti-HA antibody. Anti-TBP1 antibody was used as loading control. WT cells were included as negative control.

### atDjBs can maintain strong [*PSI*^+^] universally and weak [*PSI*^+^] differentially

As atDjBs could perform the essential functions of Sis1, we next asked if these proteins could also remodel different protein aggregates. Since very little is known about aggregate prone proteins in plants and robust aggregate remodeling assays are yet to be standardized for plant JDPs, we choose yeast prions as model protein aggregates as they have been extensively used to understand the role of molecular chaperones, especially JDPs and Hsp104 in aggregate remodeling and solubilization (45, 51).We utilized four different well studied variants of the [*PSI*^+^] prion in yeast: the strong [*PSI*^+^]^Sc4^ and [*PSI*^+^]^VH^ and the weak [*PSI*^+^]^Sc37^ and [*PSI*^+^]^VL^ variants (52, 53). Prion-bearing *sis1Δ* strains expressing *URA3-SIS1* were transformed with *CEN-TEF* plasmids expressing individual atDjB1–6 proteins. 5-FOA was again used to counterselect against the *URA3-SIS1* plasmid, and then strains expressing individual atDjB proteins were analyzed for prion maintenance. All six atDjBs were able to support strong variants of [*PSI*^+^], as confirmed by colony color assay (Fig. 6A). However, only atDjB2 and atDjB3 were able to maintain the weak [*PSI*^+^] variant [*PSI*^+^]^Sc37^, whereas no plant ortholog maintained [*PSI*^+^]^VL^. To confirm if colony color accurately reflects the aggregate status of the prion, particularly for the [*PSI*^+^]^Sc37^ strains, semi-denaturing detergent agarose gel electrophoresis (SDDAGE), a biochemical assay for resolving and visualizing detergent-resistant protein aggregates was used. Indeed, large detergent-resistant Sup35 aggregates of variable sizes, indicative of prion amyloid aggregates, were observed only in cells expressing atDjB2 and atDjB3, confirming the colony color results (Fig. 6B). The genetic background of a yeast strain sometimes impacts results of yeast prion assays (54). To rule out an impact of a particular genetic polymorphism of the specific yeast background we originally chose (W303), we repeated all of our experiments in a second genetic background (74D-694). [*PSI*^+^] maintenance results were entirely congruent between the two backgrounds (Fig. S6).

**Fig. 6.**
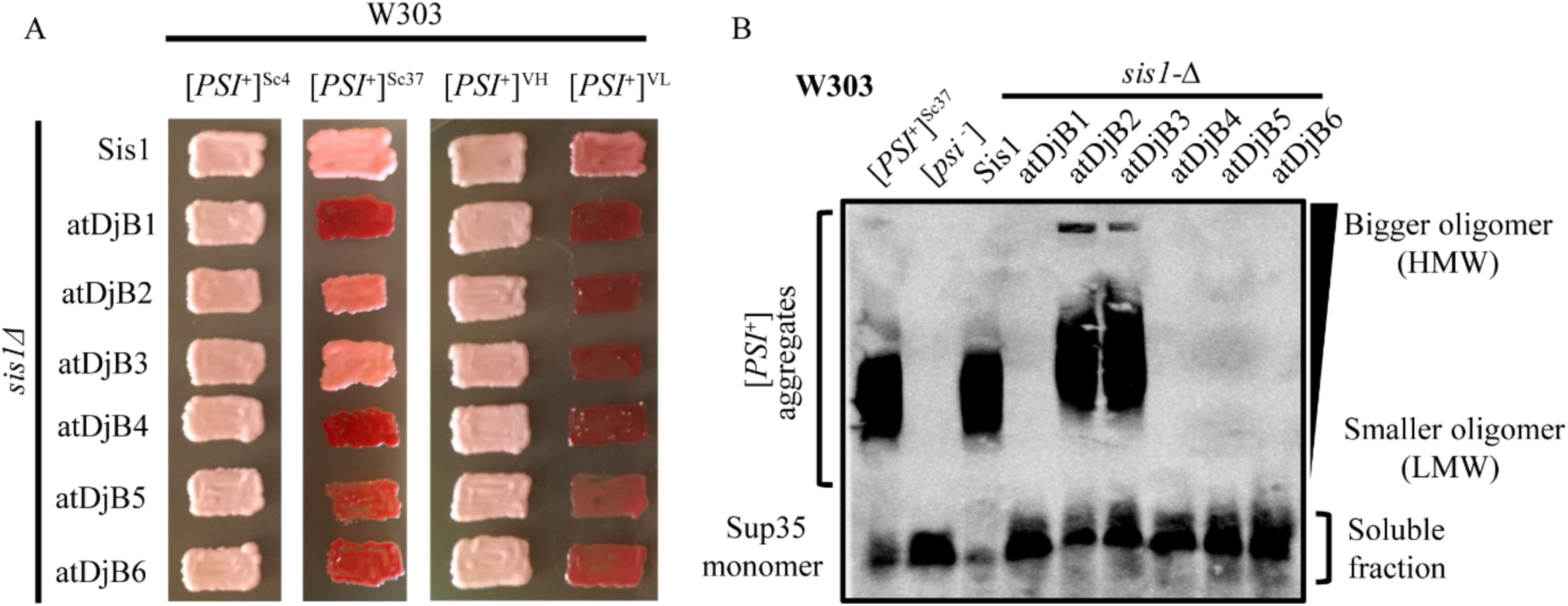
atDjBs all support strong [*PSI*^+^] but differentially maintain weak [*PSI*^+^] variants. **A.** [*PSI*^+^]^Sc4^, [*PSI*^+^]^Sc37^, [*PSI*^+^]^VH^, and [*PSI*^+^]^VL^*sis1Δ* [*URA3*-*SIS1*] cells were transformed with plasmid (pRS414) expressing Sis1 or atDjB1-6 and subjected to plasmid shuffling on 5-fluorooratic acid (5-FOA). Cells after plasmid shuffling were assayed for [*PSI*^+^] maintenance by colony color on YPD medium. Color phenotype assays are shown for representative transformants (n≥10) including parental strain for comparison. **B.** The maintenance of [*PSI*^+^]^Sc37^ was further confirmed by semi-denaturing detergent agarose gel electrophoresis (SDDAGE). Equal amount of cell lysate prepared from shuffled strains were resolved by SDDAGE, electroblotted, and probed with anti-Sup35 antibody. [*PSI*^+^]^Sc37^ and GdnHCl-treated [*psi*^-^] parent cells were included for comparison.

### atDjBs cannot substitute for Sis1 in Hsp104-mediated elimination of strong [*PSI*^+^] variants

As all six tested atDjBs equally maintained strong [*PSI*^+^] variants, we next investigated whether another Sis1 function may have been conserved among all or any plant ortholog. [*PSI*^+^], but not other prions, is efficiently eliminated from cell populations in a JDP-regulated process when Hsp104 is overexpressed (43, 55, 56). Because Sis1 is essential for the elimination of strong, but not the weak [*PSI*^+^] variant, we asked whether atDjB1–6 could substitute for Sis1. Cells bearing strong [*PSI*^+^] variants in both genetic backgrounds and expressing individual atDjB1–6 proteins in place of Sis1 were transformed with an overexpression construct of Hsp104 (*GPD-HSP104*). Selected transformants were patched on rich YPD media to check for prion maintenance. Interestingly, none of the tested atDjBs were able to substitute for Sis1 in Hsp104-mediated curing of strong [*PSI*^+^] as shown by the colony color assay (Fig. 7A). In contrast to strong variants, we recently found that the curing of weak [*PSI*^+^] variants by Hsp104 appears to be Sis1 independent (43). Accordingly, we would expect that the curing of these variants would be unaffected by the replacement of Sis1 with an atDjB protein. To test this, strains expressing atDjB2 and atDjB3 that can maintain the weak [*PSI*^+^]^Sc37^ variant were also examined for Hsp104-mediated curing. As expected, the prion was eliminated in both instances as indicated by colony color (Fig. 7B) and SDDAGE (Fig. 7C). These results suggest that plant Sis1 orthologs are deficient in the specific Sis1 functionality needed for Hsp104 to eliminate strong [*PSI*^+^] variants.

**Fig. 7.**
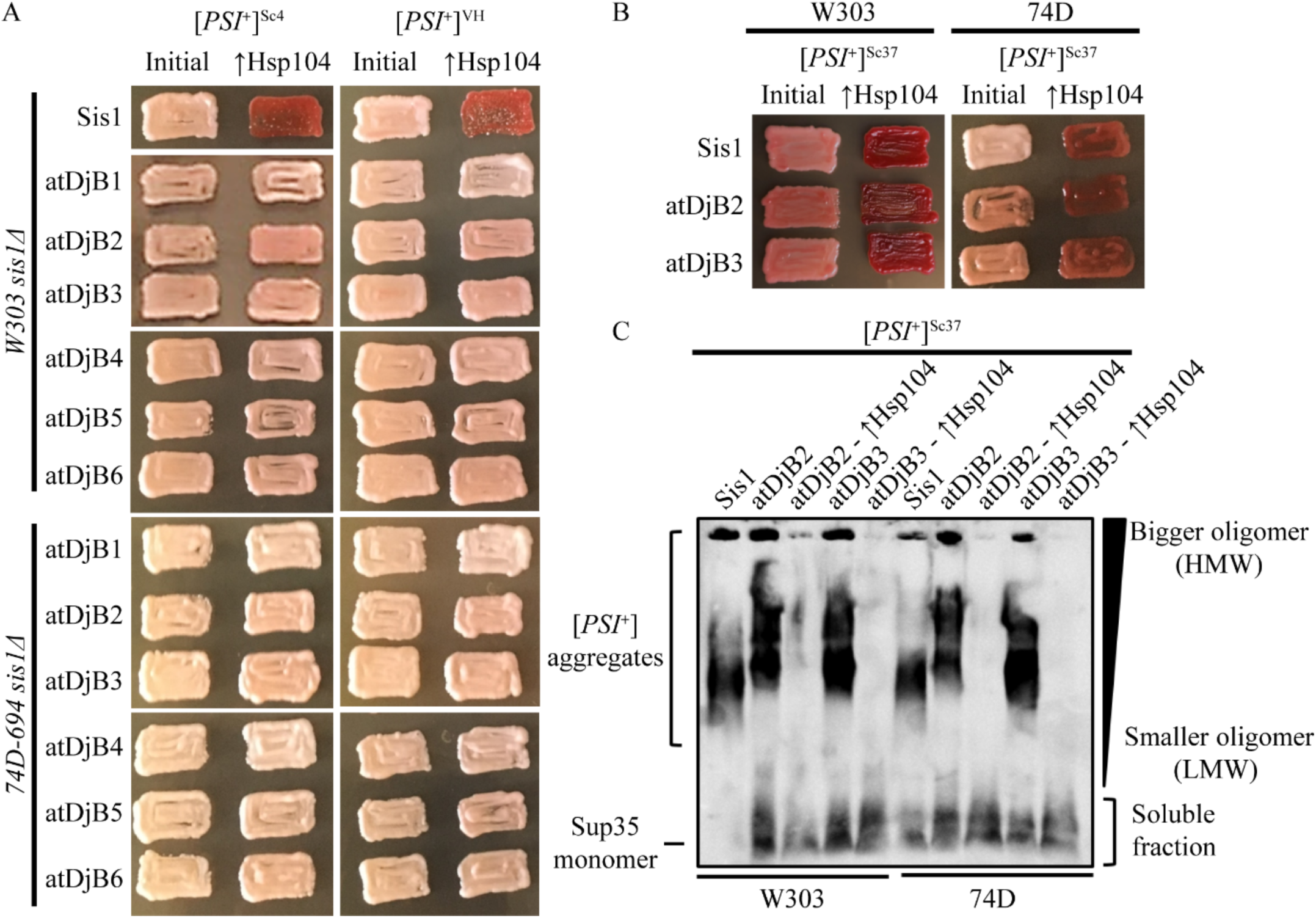
JDP requirements for Hsp104-mediated elimination of [*PSI*^+^] are prion variant dependent. **A.** [*PSI*^+^]^Sc4^ and [*PSI*^+^]^VH^ in W303 or 74D-694 *sis1Δ* shuffled strains expressing Sis1 or atDjB1-6 were transformed with a plasmid overexpressing Hsp104 (pRS426-*GPD*-*HSP104*). Cells from individual transformations were assayed for [*PSI*^+^] curing by colony color on YPD media. Color phenotype assays are shown for representative transformants (n *≥* 10).**B.** Same as panel A, but cells have [*PSI*^+^]^Sc37^ and express Sis1, atDjB2, or atDjB3. **C.** [*PSI*^+^]-status of cells shown in B is confirmed by semi-denaturing detergent agarose gel electrophoresis (SDDAGE). Equal amount of cell lysate prepared from shuffled strains from panel B, were resolved by SDDAGE, electroblotted, and probed with anti-Sup35 antibody.

### Six atDjBs differentially maintain [*RNQ*^+^] and segregate into three sets of isofunctional pairs

The observed functional diversity among atDjB1–6, revealed in the [*PSI*^+^] maintenance experiments, prompted us to ask whether these proteins can also propagate the [*RNQ*^+^] prion, as [*RNQ*^+^] propagation requires different functions of Sis1 than either strong or weak [*PSI*^+^] variants (46, 50). To examine the ability of atDjBs for maintenance of [*RNQ*^+^], we used a well-studied variant of [*RNQ*^+^] called [*RNQ*^+^]^STR^ (31). After plasmid-shuffling using 5-FOA to express individual atDjB1–6 proteins in W303 *sis1*Δ, [*RNQ*^+^]^STR^ cells, these strains were then transformed with plasmids expressing a Rnq1-GFP fusion construct. The presence of [*RNQ*^+^] in these cells can readily be detected by fluorescent foci in cells expressing Rnq1-GFP, whereas in [*rnq*^-^] cells, the fluorescence remains diffuse and homogeneously distributed throughout the cytoplasm. In contrast to strong [*PSI*^+^], atDjB1–6 differed in their ability to maintain the [*RNQ*^+^] prion, yet the pattern of maintenance was also unlike that of weak [*PSI*^+^]^Sc37^. Cells expressing atDjB1–4 as the only Sis1 ortholog, exhibited fluorescent foci like the endogenous Sis1-expressing cells, indicative of [*RNQ*^+^] maintenance. In contrast, atDjB5 and atDjB6 cells exhibited diffuse Rnq1-GFP fluorescence about the cytoplasm, indicative of [*rnq*^-^] cells (Fig. 8A), suggesting that atDjB5 and atDjB6 were unable to propagate the [*RNQ*^+^] prion in the W303 background. SDDAGE analysis using an anti-Rnq1 antibody confirmed our microscopy observations: high molecular weight [*RNQ*^+^] aggregates were observed only in cells expressing atDjB1–4. Cells expressing either atDjB5 or atDjB6 exhibited only the monomeric form of Rnq1, indicating that they are [*rnq*^-^] (Fig. 8B). Once again, we repeated these experiments using SDDAGE in the 74D-694 background with identical results (Fig. S7). Put together, our results show that the six tested Sis1 orthologs may have distinct aggregate remodeling functionalities in *A thaliana*.

**Fig. 8.**
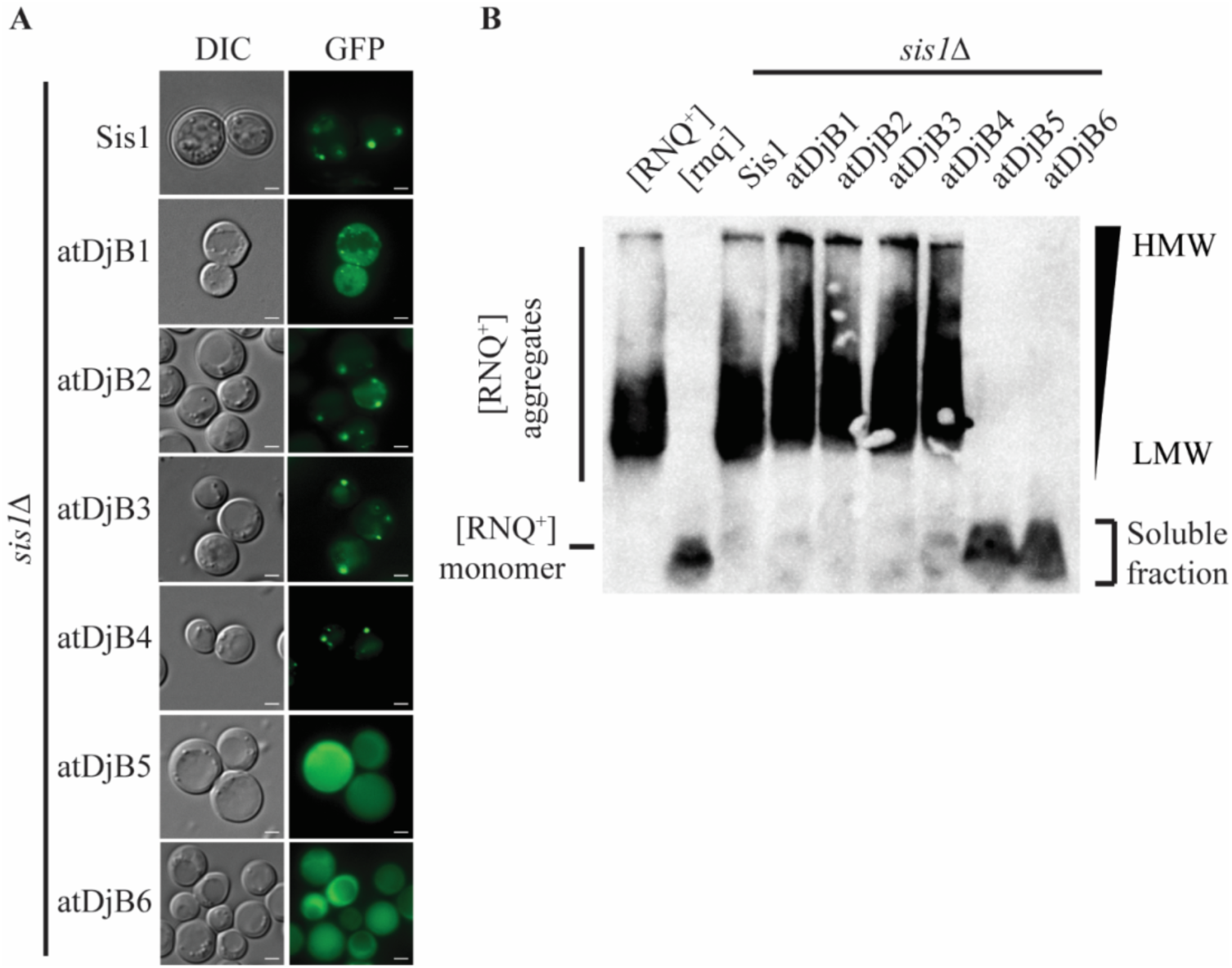
Maintenance of [*RNQ*^+^] by atDjBs. **A.** [*RNQ*^+^] *sis1Δ* [*URA3*-*SIS1*] cells were transformed with plasmid (pRS414) expressing Sis1 or atDjB1-6, subjected to plasmid shuffling on 5-fluorooratic acid (5-FOA), and then transformed with a Rnq1-GFP reporter plasmid (pRS413-*TEF*-*RNQ1*-GFP) followed by fluorescence microscopy analysis. Scale bar, 2 μm. **B.** Equal amounts of cell lysate prepared from shuffled strains from panel **A** were resolved on semi-denaturing detergent agarose gel electrophoresis (SDDAGE), electroblotted, and probed with anti-Rnq1 antibody. Control [*RNQ*^+^] and GdnHCl-treated [*rnq*^-^] parent cells were included for comparison.

### atDjB1–6 differentially substitute for Ydj1 in budding yeast

Sis1 shares significant functional overlap with Ydj1, the major Class I JDP of yeast (57). The C-terminal domains of one or the other protein are required for the viability of yeast cells (58). Ydj1 performs several housekeeping chaperone functions and is important for growth at high temperatures (59). *ydj1Δ* cells grow slowly at 25°C and are dead at 34°C. The slow growth phenotype of *ydj1Δ* is rescued by overexpressing Sis1 (60). To test for another potentially-conserved function of Sis1, specifically one that is dependent on Sis1’s C-terminal domain, we asked whether atDjB1–6 can also rescue the slow growth phenotype of *ydj1Δ*. To address this, *ydj1Δ* cells were transformed with plasmids expressing individual atDjBs. Interestingly, while all these JDPs rescued the essential functions of Sis1, only atDjB1–4 could rescue the slow growth of the *ydj1Δ* strain (Fig. 9A). To more rigorously assess the functional differences within atDjB1–6, we tested their ability to rescue the lethality of *sis1Δydj1Δ* following the loss of [*URA3-SIS1*]. As expected, only atDjB1–4, could substitute for Sis1 deficiency in the absence of Ydj1. Previously, it has been shown that a Sis1^1-121^ fragment (which lacks the C-terminal domains) is sufficient to rescue *sis1Δ*cells but only in the presence of full-length Ydj1 (58). Plasmids expressing different atDjBs were first transformed into *sis1Δydj1Δ* [*URA3-SIS1*] pRS313-Sis1^1-121^ cells and then counterselected against the *URA3-SIS1* plasmid on 5-FOA plates. We found that atDjB1–4 rescued growth of *sis1Δydj1Δ* pRS313-Sis1^1-121^ cells like Sis1 at 30°C, but, again, atDjB5 and atDjB6 did not (Fig. 9B; 9C). These results indicate that atDjB5 and atDjB6 are functionally distinct from other atDjBs.

**Fig. 9.**
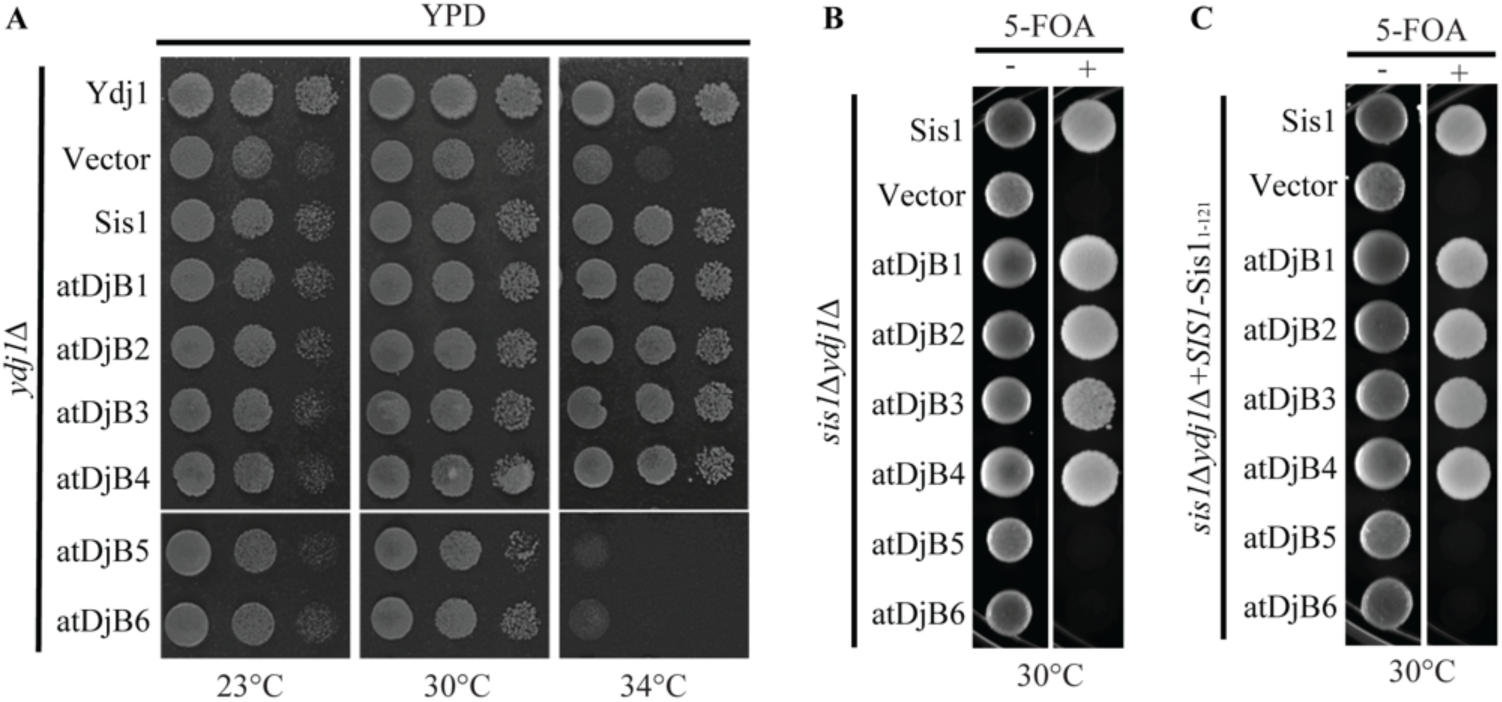
atDjB1-6 differentially rescue the generalized functions of Ydj1 in *S. cerevisiae*. **A.** Equal volume of ten-fold serial dilutions of *ydj1Δ* cells harboring empty pRS414 plasmid (vector) or pRS414 expressing Ydj1, Sis1 or atDjB1-6 were spotted on YPD plates and incubated at 23°C, 30°C and 34°C for 3 days. **B.** Equal volume of ten-fold serial dilutions of *sis1Δydj1Δ* [*URA3*-*SIS1*] harboring empty pRS414 plasmid (vector) or pRS414 expressing Sis1 or atDjB1-6were spotted on media with (+) and without (-) 5-fluoroorotic acid (5-FOA) and incubated at 30°C for 4 days. **C.** Equal volume of ten-fold serial dilutions of *sis1Δydj1Δ* [*URA3*-*SIS1* and pRS313-Sis1_1-121_] harboring empty pRS414 plasmid (vector) and pRS414 expressing Sis1 or atDjB1-6were spotted on media with (+) and without (-) 5-fluoroorotic acid (5-FOA) and incubated at 30°C for 4 days.

### The glycine-rich regions determine the functional specificity of atDjBs toward the [*RNQ*^+^] prion

Since all the atDjBs rescued the essential functions of Sis1, we hypothesized that the functional differences observed in atDjB5 and atDjB6, including their inability to maintain [*RNQ*^+^] prion, may be due to differences between the glycine-rich regions of these proteins and these same regions in other orthologs which retain more Sis1 functions, like atDjB2.Previous reports have also demonstrated that the glycine-rich region of Sis1 plays a critical role in the propagation of [*RNQ*^+^] (32, 61). Additionally, atDjB2 and atDjB5 have significant differences in their glycine-rich regions (Fig. S8). As a first step toward identifying the specific sequence features that give rise to the distinct functionalities conserved among some but not other atDjBs, we asked whether the glycine-rich region can direct atDjB chaperone action. To do this, we made two chimeras of atDjB5 with either the glycine/phenylalanine and glycine/methionine (G/F and G/M) (Chimera A) or only the G/F (Chimera B) region(s) of atDjB2 (Fig. 10A). These chimeras allowed us to ask whether sequence differences in the glycine-rich region are contributing to the differential aggregate remodeling properties among the Arabidopsis atDjBs.

**Fig. 10.**
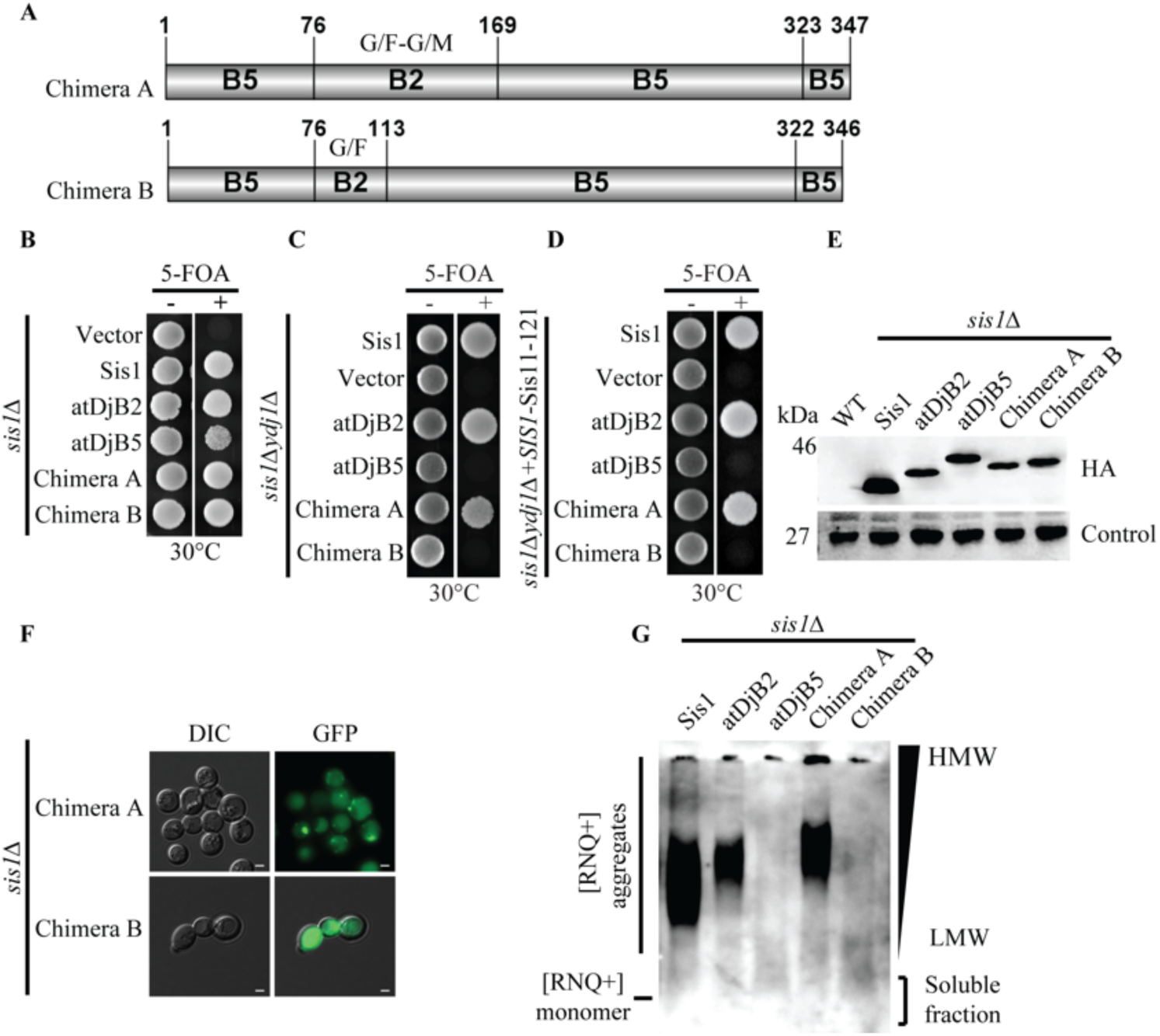
Conserved GF/GM region defines the functional specificity among atDjBs. **A.** Domain organization of Chimera A and B. Chimera A contains the following amino acids: B5 (1-76):B2 (80-173):B5 (172-349). Chimera B contains the following amino acids: B5 (1-76):B2 (80-117):B5 (118-349). Please see the MATERIALS AND METHODS for details about the construction of Chimera A and Chimera B. **B.** Equal volume of ten-fold serial dilutions of *sis1Δ* [*URA3*-*SIS1*] cells harboring empty pRS414 plasmid (vector) or pRS414 expressing Sis1 or Chimera A/B were spotted on media with (+) or without (-) 5-fluoroorotic acid (5-FOA) and incubated at 30°C for 3 days. **C.** Equal volume of ten-fold serial dilutions of *sis1Δydj1Δ* [*URA3*-*SIS1*] harboring empty pRS414 plasmid (vector) and pRS414 expressing Sis1 or Chimera A/B were spotted on media with (+) and without (-) 5-fluoroorotic acid (5-FOA) and incubated at 30°C for 4 days. **D.** Equal volume of ten-fold serial dilutions of *sis1Δydj1Δ* [*URA3*-*SIS1* and pRS313-Sis1^1-121^] harboring empty pRS414 plasmid (vector) and pRS414 expressing Sis1 or Chimera A/B were spotted on media with (+) and without (-) 5-fluoroorotic acid (5-FOA) and incubated at 30°C for 4 days. **E.** Equal amounts of total cell lysate prepared from *sis1Δ* cells harboring plasmids expressing HA-tagged constructs of Sis1 and Chimera A/B were resolved on SDS-PAGE, electroblotted, and probed with anti-HA antibody. Anti-TBP1 antibody was used as loading control. WT cells were included as negative control. **F.** *sis1Δ* cells [*URA3*-*SIS1*] bearing [*RNQ*^+^] examined in this study was used for subsequent transformation of each shuffled strain by a Rnq1-GFP reporter plasmid (pRS413-*TEF*-*RNQ1-GFP*) followed by fluorescence microscopy analysis. Scale bar, 2 μm. **G.** Protein lysate prepared from shuffled strain of *sis1Δ* cells harboring constructs of Sis1 or Chimera A/B were resolved on semi-denaturing detergent agarose gel electrophoresis (SDDAGE), electroblotted, and probed with anti-Rnq1 antibody.

Before analyzing the aggregate remodeling properties of the chimeras, we verified if they behaved like atDjB2 in growth assays by testing the ability of Chimeras A and B to rescue the lethality of *sis1Δ*, *sis1Δydj1Δ*, or *sis1Δydj1Δ* pRS313-Sis1^1-121^ strains. While Chimera A could substitute for Sis1 deficiency in all strains, Chimera B supported the growth of only *sis1Δ* but not of *sis1Δydj1Δ* or *sis1Δydj1Δ* pRS313-Sis1^1-121^double deletion strains (Fig. 10B; 10C; 10D). HA-tagging followed by western analysis revealed that these proteins are expressed at similar levels in yeast cells, so this functional difference is likely not due to differences in protein quantities (Fig. 10E). Next, we determined the ability of the chimeras to propagate [*RNQ*^+^]. *sis1Δ* [*RNQ*^+^] cells expressing atDjB2, atDjB5, Chimera A, or Chimera B were transformed with a plasmid expressing Rnq1-GFP. Chimera A maintained [*RNQ*^+^] like atDjB2 as indicated by distinct fluorescent foci (Fig. 10F). However, Chimera B, which only has the GF region of atDjB2, was unable to propagate [*RNQ*^+^] (Fig. 10F). These results were also confirmed by SDDAGE (Fig. 10G). Finally, we asked whether the glycine-rich regions of atDjB2 also conferred the ability to propagate weak [*PSI*^+^]^Sc37^ to the atDjB5 protein. Consistent with the previous results for atDjB5, both chimeras were only able to support the strong but not the weak [*PSI*^+^] variants, as confirmed by colony color assay (Fig. S9A) and SDDAGE (Fig. S9B) in W303 cells and in the 74D-694 background (Fig. S9C; S9D) as well. Collectively, these results indicate that the biochemical differences among atDjBs that allow the rescue of *sis1Δydj1Δ* strains, and propagation of the [*RNQ*^+^] prion, are specifically determined by the glycine-rich regions of these proteins.

## Discussion

Here we report the identification and characterization of aggregate remodelling Class II JDPs (atDjBs) in *A. thaliana*. Results presented in this study demonstrate that atDjBs interact with heat-induced protein aggregates and co-localize with the major disaggregase Hsp101, at distinct protein aggregate centers (PACs) in plant cells. The fact that these proteins displayed specificity towards different types of aggregates suggests that plants may employ different JDPs to specify and modulate the aggregate remodelling activities of the Hsp70-Hsp100 bi-chaperone system.

Formation of cytotoxic protein aggregates is the hallmark of cellular stress, particularly, high temperature (62, 63). Additionally, environmental stresses like UV exposure, low temperature, and high salt also induce protein misfolding and aggregation in plant cells (64, 65). Perhaps unsurprisingly, while several chaperones are heat-shock regulated (66), some are either constitutively expressed or regulated by salt, cold, osmotic, and oxidative stress (67). atDjBs studied here exhibited variability in their expression profiles during different developmental stages and stress conditions suggesting that besides mitigating protein misfolding and aggregation under stress conditions, atDjBs may also play important roles in plant growth and development under non-stress regimes.

Specialized JDPs recruit Hsp70 to the aggregates and concomitantly stimulate their weak ATPase activity (68). Hsp70 then recruits and activates Hsp100 to extract proteins from a variety of protein aggregates, a crucial step in aggregate remodeling (69, 70). A Similar, yet complex scenario emerges in plants. While atDjB1-6 co-localized with Hsp101 to distinct heat-induced PACs in isolated Arabidopsis protoplasts, this might not be physiological, as out of the six JDPs analyzed here, only atDjB2 and atDjB3 were heat-inducible. Because Hsp101 is also heat-inducible, it is plausible that among the six atDjBs, only atDjB2 and atDjB3 play specialized roles in regulating the disaggrergase activity of Hsp70-Hsp100 chaperone machines in response to high temperature. At this point we cannot rule out the possibility that atDjB1, atDjB4, atDjB5 and atDjB6 cooperate with Hsp101 *in vivo*. It is possible that these atDjBs drive the disaggregase activity of Hsp70s without Hsp100. Such standalone aggregate remodeling systems are known to exist in metazoans which lack a Hsp100 homolog (except in the mitochondria) (71).

All the atDjBs we fully analyzed here rescued the essential functions of Sis1 in a *sis1Δ* strain. This requires the Hsp70 co-chaperone activity, as mutations in the critical HPD motif within the J-domain of atDjB1 abolishes their ability to rescue a *sis1Δ* strain (44). Besides performing an unknown essential function, Sis1 also remodels different types of protein aggregates, including prions in yeast (35, 72, 73). Distinct domains of Sis1 are important for the maintenance of different yeast prions (31, 43). Interestingly, atDjB1-6 exhibited different aggregate remodeling properties in yeast, suggesting towards a subfunctionalization of the aggregate remodeling activities within this set of proteins. Except for the glycine-rich (G/F-G/M) region following the J-domain, atDjBs show significant similarity with each other as well as with their yeast homolog Sis1. The G/F region of Sis1 is required for [*RNQ*^+^] maintenance (32). Moreover, this region shares a functional overlap with the G/M region and carries determinants for Sis1’s specificity (61). Consistent with this, results from protein chimera experiments indicate that variations in the glycine-rich region of atDjBs not only defines their functionality but also their specificity towards different types of aggregates. Most notably, replacement of the G/F+G/M-rich region of atDjB5, which cannot maintain [*RNQ*^+^], with those from atDjB2 imparted the ability to propagate [*RNQ*^+^]. However, the replacement of only the G/F-rich region was insufficient, indicating that both glycine-rich regions are required. The fact that the chimera with both glycine-rich regions substituted, rescued the lethality of *sis1Δydj1Δ* and *sis1Δydj1Δ* pRS313-Sis1^1-121^ strains also implies that the glycine-rich regions in at least some atDjBs may have functional redundancy with their C-terminal client binding domains. This is true for Sis1 as well. While complete deletion of the G/F region results invariantly in the loss of [*RNQ*^+^], point mutations in the unique region (amino acids 101-113) in the G/F result in [*RNQ*^+^] loss only in the absence of Sis1’s C-terminal domains (61). Interestingly, despite substituting for Sis1 in all other functions, the fully-swapped G/F and G/M chimera did not propagate weak [*PSI*^+^]^Sc37^, indicating that some sort of cooperativity between the glycine rich region and the CTD, possibly to mediate a bipartite interaction with Hsp70 (74)may be necessary for the remodeling of some aggregate.

Overexpression of Hsp104 is known to rapidly cure strong variants of [*PSI*^+^] from cell populations by a highly-debated mechanism, requiring Sis1 (54). Strikingly, all six orthologs examined were deficient in replacing Sis1 in this process which is congruent with previous findings for both Hdj1 and Droj1 (43, 54). Thus, regarding Hsp104-overexpression mediated aggregate remodeling resulting into prion loss, all 8 higher eukaryotic Sis1 orthologs that have been studied specifically lack this ability. These eukaryotic orthologs, together with Sis1, now form an exciting set of sequence-similar proteins with diverse but clearly defined aggregate remodeling functionalities. While no atDjB behaved exactly like Sis1, our data resolved the six orthologs into 3 sets of isofunctional pairs: atDjB2/3 maintained almost all prions tested, most similar to Sis1; atDjB1/4 maintained [*RNQ*^+^] and strong [*PSI*^+^]variants but not [*PSI*^+^]^Sc37^, like the human and Drosophila orthologs Hdj1 and Droj1 (31, 43); and atDjB5/6 fail to maintain any prion except strong [*PSI*^+^] variants, similar to the Sis1 mutant that disrupts the bipartite interaction between Sis1 and Hsp70 (75).

Put together, this study demonstrates that compared to bacteria and budding yeast which have one each (CbpA and Sis1 respectively), or even complex metazoans like humans which have three (DNAJB1, DNAJB4 and DNAJB5) (68), Sis1-like Class II JDPs have proliferated in plants with seven in Rice and potato, nine in maize (*unpublished results*) and eight in *A. thaliana* (76), which could result in a highly complex network of aggregate-remodeling chaperone systems. There is a growing list of plant proteins that show amyloid behaviour. This includes the naturally occurring monellin in *Dioscoreophyllum cumminsii*, transglutaminase (TGZ) from maize, seed storage proteins in pea, soybean and wheat, defensin protein in radish and coconut, and prohevein protein from rubber tree (77–86). Recently flowering pathway proteins, Luminidependens (LD), Flowering Locus PA (FPA) and Flowering Locus CA (FCA) from *A. thaliana* were shown to behave like prions in yeast cells (87). Additionally, studies show that several plant proteins harbour amino acid clusters that are prone to aggregate (87, 88). Some potential prion-forming proteins may have roles in regulating different aspects of plant growth and development (87, 89). While the present study identifies stress associated JDPs that might collaborate with Hsp101 to solubilize heat-induced protein aggregates, it remains to be seen if they also remodel different amyloid aggregates thereby acting as epigenetic modifiers of plant growth and development in Arabidopsis.

## Experimental procedures

### In silico analysis

atDjB domain architecture was predicted by SMART database (http://smart.embl-heidelberg.de/) (90). Models for domain organization were created by IBS (http://ibs.biocuckoo.org/index.php) (91). Secondary structures were predicted using SWISS-MODEL (https://swissmodel.expasy.org) (92). EMBOSS needle was used for the identification of sequence similarity and identity between Sis1 orthologs (https://www.ebi.ac.uk/Tools/psa/emboss_needle/) (93).

### Plasmid construction for yeast complementation

Open reading frames (ORFs) corresponding to full-length Sis1 orthologs were PCR amplified from a pooled *A. thaliana* cDNA made from RNA isolated from stressed and unstressed shoots, roots, and inflorescence. *S. cerevisiae SIS1* was amplified from yeast genomic DNA. atDjB1, atDjB2, atDjB3, atDjB4, atDjB5 and atDjB6 encoding genes were cloned either into *HIS3*-marked pRS413 or *TRP1*-marked pRS414 yeast expression vectors under the *TEF1* promoter (50) in *Bam-HI* and *Sal-I* sites. To generate N-terminus HA-tagged constructs, the tag was added in the forward primers before the ORF of *Sc*Sis1 in between *Spe-I* and *Bam-HI* sites and cloned into *HIS3*-marked pRS413 using the same reverse primer having *Sal-I* site. ORFs of the orthologs were subcloned in the same HA-tagged construct by releasing *Sc*Sis1 fragment with *Bam-HI* and *Sal-I* sites. All the constructs were confirmed by restriction digestion and sequencing.

Overlapping PCR method was used to construct Chimera A and Chimera B from atDjB2 and atDjB5 genes. A chimeric gene encoding the J-domain (aa 1 to 76) of atDjB5, the G/F and G/M regions (aa 81 to 173) of atDjB2 and the C-terminal domains (aa 173 to 350) of atDjB5 was referred to as Chimera A. A chimeric gene encoding the J-domain (aa 1 to 76) of atDjB5, the G/F region (aa 81 to 117) of atDjB2, and part of the glycine-rich region and C-terminal domains (aa 118 to 350) of atDjB5 was called Chimera B. These were cloned into *TRP1*-marked pRS414 yeast expression vectors under the *TEF1* promoter with *Bam-HI* and *Sal-I* sites. To compare the expression levels of the Chimeric proteins, they were subcloned in previously used pRS413-HA-Sis1 construct having N-terminal HA-tag by releasing Sis1 fragment with *Bam-HI* and *Sal-I* sites.

Protein fusion constructs were generated for subcellular localization analysis, the open reading frames having *attB1* and *attB2* sites for atDjB1, atDjB2, atDjB3, atDjB4, atDjB5, and atDjB6 without a stop codon and Hsp101 with a stop codon were amplified from Arabidopsis cDNA. PCR fragments were then cloned into Gateway pDONR207 vector (Invitrogen, Thermo Fisher Scientific) by BP reaction to produce entry clones. The entry clones were confirmed through sequencing. atDjB1–6 pDONR207 entry clones were used in LR reaction with the Gateway-compatible plant binary vector pGWB5 (94) containing the 35S promoter and the C-GFP fragment to produce destination clones. Similarly, the Hsp101 pDONR207 entry clone was used in LR reaction with Gateway-compatible plant binary vector pGWB661 (95)containing the 35S promoter and the N-RFP fragment to produce destination clones. The BP and LR reactions were done as described in manufacturer’s manual (Invitrogen, Thermo Fisher Scientific). All primer sequences are listed in supporting Table 1.

### Protein purification and Antibody generation

atDjB1-C-term (aa 157 to 335) was amplified from pRS413-atDjB1 and cloned into pET28a(+) bacterial expression vector using *Bam-HI* and *Xho-I* sites. Protein was expressed from *E. coli* strain (Rosetta) and purified using Ni-NTA affinity chromatography according to the manufacturer’s instructions (Qiagen Cat No./ID: 30210). Purified protein was used as an immunogen to raise polyclonal antibody in rabbits at a local commercial facility (DPL, Bhopal, India)

### Yeast methods

Haploid *S. cerevisiae* W303 or 74D-694 derived *ydj1Δ* (*ydj1::HIS3*), *sis1Δ (sis1::LEU2)* and *sis1Δydj1Δ*(*sis1::LEU2, ydj1::HIS3*) strains previously described (44) were used throughout the study. Yeast strains with *sis1Δ*harbored a *URA3*-marked plasmid expressing the wild-type *SIS1* gene (pRS316-*SIS1*). For testing the ability of atDjBs to complement the *in vivo* functions of Sis1, we used 5-fluoroorotic acid (5-FOA) plasmid shuffling (96). atDjB1–6 cloned in yeast expression vectors were transformed into W303 or 74D-694 *sis1Δ* and*sis1Δydj1Δ*strains and counterselection of pRS316-*SIS1* was done on medium containing 5-FOA. Yeast strains bearing [*PSI*^+^]^Sc4^, [*PSI*^+^]^Sc37^, [*PSI*^+^]^VL^, [*PSI*^+^]^VH^, or [*RNQ*^+^]^STR^ prions were used to study the aggregate remodeling properties of atDjB1–6through plasmid shuffling as described previously (31). [*PSI*^+^] maintenance was observed through the colony color assay. Briefly, all the selected transformants after counterselection were patched on YPD for prion maintenance. If Sup35 is soluble or cells are in [*psi*^-^] state, the colonies look red due to accumulation of red intermediate resulting from the block in adenine biosynthesis. When Sup35 is in the aggregated state, *i.e*., [*PSI*^+^], the yeast colonies look white or pink due to complete or partial restoration of adenine biosynthesis (39). [*RNQ*^+^] maintenance in cells were observed directly under Zeiss, Apotome at Excitation/Emission ∼361/497 nm at 100X magnification following transformation by (pRS413*-*Rnq1-GFP) plasmid (37). To create [*prion*^-^] strains or to confirm prion maintenance, prion-bearing cells were treated with the Hsp104 inhibitor guanidine hydrochloride (3mM) in YPD liquid medium which readily cures [*PSI*^+^] and [*RNQ*^+^]. For examining prion loss or curing by Hsp104 overexpression, a plasmid overexpressing Hsp104 was transformed in all the prion bearing strains and incubated on selective medium to detect prion maintenance as described previously (43).

### Preparation of Soluble and Insoluble Protein Fractions

Fractionation of soluble and heat-denatured proteins was done as described previously (27). Briefly, two-week-old Arabidopsis seedlings grown on 1/2 MS plates were either untreated (control) or heat stressed as described above. For each sample, 0.7 g of plant tissue was harvested, and a crude protein extract was prepared using 1 mL of protein isolation buffer (25 mM HEPES, pH 7.5, 200 mM NaCl, 0.5 mM EDTA, 0.1% (v/v) Triton X-100, 5 mM ε-amino-*N*-caproic acid, and 1 mM benzamidine-HCl). After grinding the sample using a mortar and pestle, samples were further homogenized with a Dounce glass tissue grinder for 1 min on ice. The crude protein extract was transferred to a microcentrifuge tube, and 500 μL was used for separation into soluble and insoluble fractions, while 300 μL was added to 100 μL of 4X SDS-PAGE sample buffer (8% [w/v] SDS, 46% [v/v] glycerol, 20% [v/v] β-mercaptoethanol, 250 mM Tris, pH 6.8, and 0.01% [w/v] Bromophenol Blue). For fractionation, samples were spun in a table top centrifuge at 16,100*g* for 15 min. The supernatant was collected in a fresh microcentrifuge tube, and 300 μL of supernatant was added to 100 μL of 4× SDS-PAGE sample buffer for SDS-PAGE and immunoblot analysis. To facilitate washing of the insoluble fraction, 0.1 g of quartz salt (Sigma-Aldrich) was added to the pellet fraction, and samples were washed seven times with 1 mL of protein isolation buffer. For each wash, the pellet was resuspended by pipetting and intermittent mixing, and the samples were centrifuged subsequently at 16,100*g* for 15 min. After the washes, the insoluble fraction was washed once with PIB without Triton-X100. The pellet fraction was resuspended in 250μL of PIB and along with equal volume of 2X SDS-PAGE sample buffer, creating a total volume of 500 μL. Samples were spun for 30 s at 1,500*g*, and the soluble fraction was transferred to a new microcentrifuge tube.

### Protein Analyses

Total protein from yeast was isolated from cultures in log phase by treating cells with 0.1 N NaOH and resuspending in SDS sample buffer (62.5 mM Tris·HCl, pH 6.8, 5% glycerol, 2% SDS, 2% β-mercaptoethanol, and 0.01% bromophenol blue). Proteins were resolved on 12% SDS-PAGE, electroblotted on PVDF membrane (Bio-Rad), and for immunodetection probed with anti-HA antibody (Sigma-Aldrich) or an anti-TBP antibody (a kind gift from Prof R. S. Tomar, IISER Bhopal, India).

Semi-denaturing detergent agarose gel electrophoresis (SDDAGE), a method for resolving detergent-resistant aggregates, was used to confirm the presence of [*RNQ*^+^] and [*PSI*^+^] as previously described (31). Briefly, cells were lysed by vortexing at 4°C with sterile glass beads. Following centrifugation at 4°C, cleared lysates were mixed with SDS loading buffer and incubated at 25°C for 7-8 minutes. Aggregates were resolved in a 1% agarose gel made in 1.5% (^w^/^v^) Tris-glycine and 0.1 %SDS (SeaKem Gold PFGE agarose). Proteins were transferred to a nitrocellulose membrane at 1A for 1 h in a Tris-glycine/methanol buffer. Prion aggregates were visualized by performing western analysis using antibodies specific for either Rnq1 or Sup35 (gifts from the Craig and Tuite labs, respectively).

Total, soluble and pellet plant protein fractions were separated by SDS-PAGE on 13% SDS-PAGE, electroblotted on PVDF membrane (Bio-Rad), and processed for immunodetection. Primary antibody dilutions and Agrisera order numbers were as follows: CI sHSPs (1:3,000; AS07 254); GAPC (1:1000; AS15 2894), and C-term atDjB1 (generated in the laboratory; 1:1,000). Blots prepared for fluorescent detection were incubated with goat anti-rabbit secondary antibody, Alexa Fluor Plus 680 (1:20,000; A32734-Invitrogen), and detected using a Li-Cor Odyssey.

### Plant assays

The Col-0 ecotype of *Arabidopsis thaliana* was used in all experiments. Seeds were surface sterilized and grown on plates with 1/2MS Basal Salt Mixture (M5524: Sigma), MES (0.05%) [(2-(N-morpholino) ethane sulfonic acid], 1% (w/v) sucrose, and 0.7% agar (PCT0901: HIMEDIA), stratified at 4°C for 48 h in the dark and then transferred to a Percival LED22C8 growth cabinet at 22°C Day/18°C Night and 70% humidity (light intensity 120±20 μmol m^−2^ s^−1^, 16-h light:8-h dark cycle). After germination, 20-25 days plantlets were subjected to various stress treatments (heat stress by 37°C for 1h, cold stress by 4°C for 24h, mannitol (150mM) for 24h, salt stress (150mM NaCl) for 24h, and mechanical stress by puncturing the leaves and leaving them on media for 1h).

For protoplast isolation 15-day old Arabidopsis seedlings were transferred to the soil and grown under short day conditions in a Percival LED22C8 growth cabinet at 22°C Day/18°C Night and 70% humidity (light intensity 120±20 μmol m^−2^ s^−1^, 10-h light:14-h dark cycle). 3 to 4-week-old plants were used for protoplast isolation and transfection.

The atDjB1-B6 C-GFP fusion constructs (pGDW5) were transformed into protoplasts by using polyethylene glycol-mediated transformation as described previously (97). Hsp101 N-RFP fusion construct (pGWB661) was used as protein aggregate centres (PAC) marker for co-localization studies. Images were taken with an Olympus FluoView-3000 confocal inverted laser scanning confocal imaging system (Olympus, Japan). Excitation and emission wavelengths Ex488/Em500-520, Ex558/Em580-600, and Ex594/Em630-700 were used for GFP, RFP and Cy5, respectively. The images acquired from the confocal microscope were analyzed using ImageJ software (98). Separate channel images were assembled using Adobe Photoshop CS5.1.

### RNA extraction and quantitative reverse transcription PCR

Total RNA was extracted from plant seedling samples using TRizol reagent (SIGMA) and reverse transcribed to make cDNA after DNase I treatment using iScript cDNA synthesis kit (Bio-Rad) following the manufacturer’s instructions. Real-time PCR was performed in a CFX connect 96-well real-time system using iTaq universal SyBr green Supermix (BioRad). Data analysis was done according to MIQE guidelines (99). Reference gene used was *ACTIN* for normalization.

### Data availability

All the data reported is in the manuscript.

## Acknowledgments

We thank Elizabeth Craig (University of Wisconsin–Madison), Mick Tuite (University of Kent), and R. S. Tomar (IISER Bhopal, India) for yeast strains, plasmids, and antibodies, Y.T. thanks the Ministry of Science and Technology for Council of Scientific and Industrial Research (CSIR) fellowship, S.S.L. thanks the Indian Ministry of Human Resource Development for a Graduate Aptitude Test in Engineering fellowship. We are grateful to Fund for Improvement of Science &Technology Infrastructure in Higher Educational Institutions (FIST) for providing live cell imaging system to the IISER Bhopal Microscopy Central Facility.

## Funding and additional information

This work was supported by funds from the Science and Engineering Research Board (EMR/ 2015/001213), intramural funds from IISER Bhopal to C.S.; the Lafayette College Chemistry Department, the EXCEL research scholarship program, the Camille and Henry Dreyfus Foundation (Award No. TH-18-017), and the National Institute of General Medical Sciences of the National Institutes of Health (Award No. R15GM110606), awarded to J.K.H. The content is solely the responsibility of the authors and does not necessarily represent the official views of the National Institutes of Health.

---

**Fig. S1.**
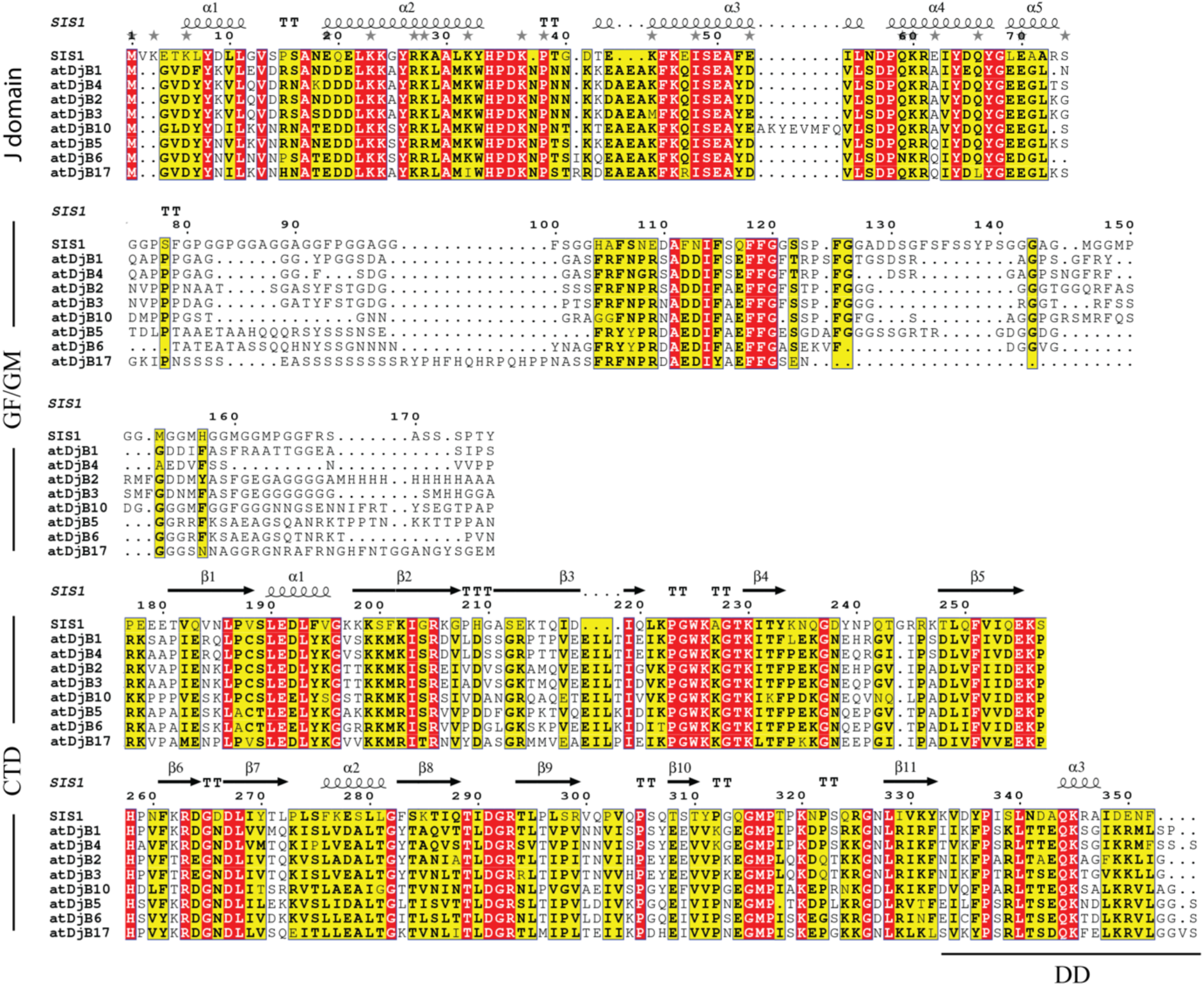
Sequence alignment of Sis1 and atDjBs. Homology comparison of amino acid sequences of Sis1 and its *A. Thaliana* atDjBs orthologs. The multiple sequence alignment was carried out using MAFFT ver.7 (100), followed by ESPript 3.0 (101). Domains are indicated. The secondary structure element for J-domain (PDB: 4RWU) and CTD (PDB: 1C3G) of Sis1 are given above their corresponding sequences. The conserved residues are indicated by the default colouring scheme of the ESPript program. Arrow, spring, TT and TTT above the sequences represent *β*-strand, helical, strict *β*-turn and strict α-turn conformations, respectively.

**Fig. S2.**
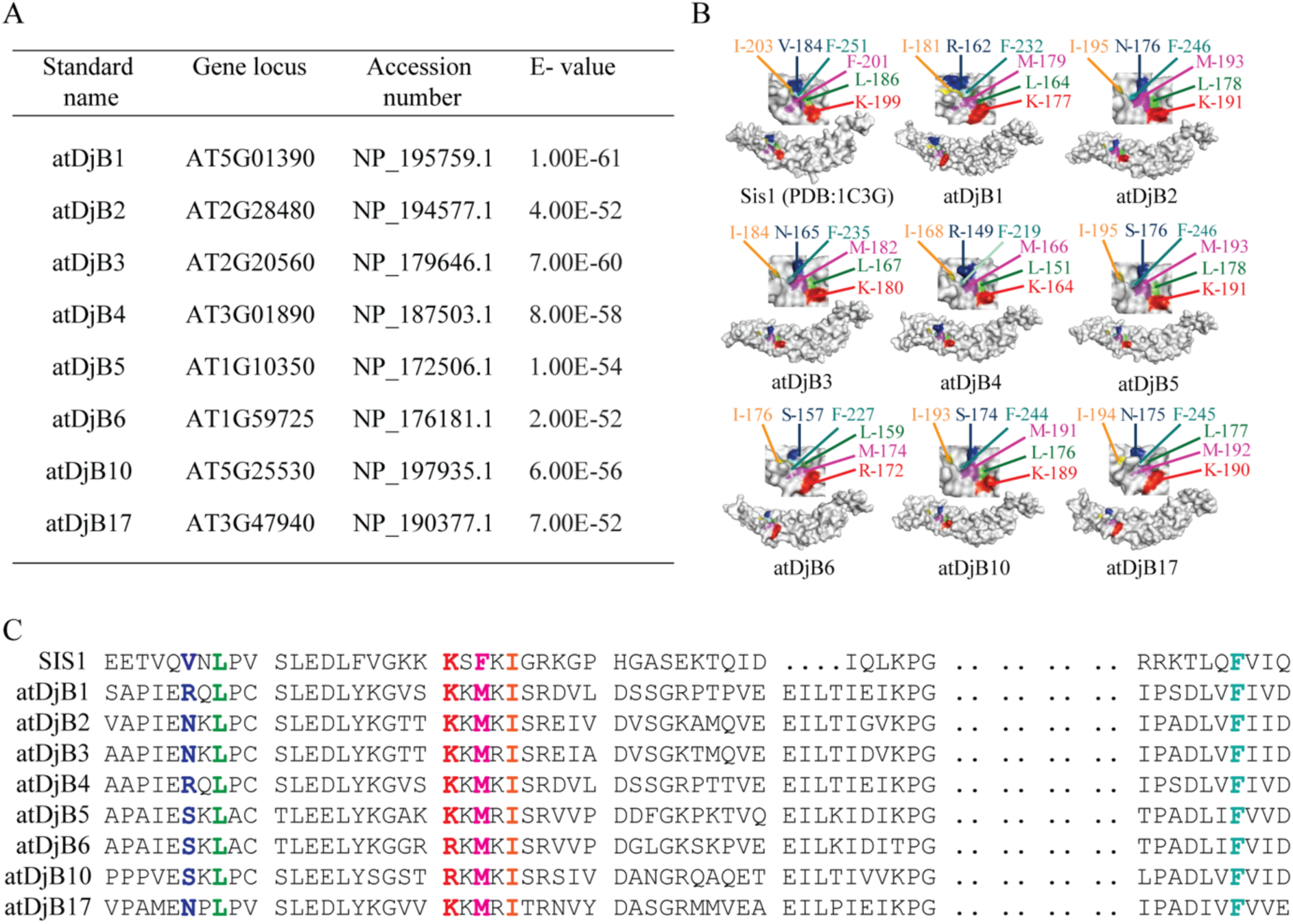
Conserved features among *A. thaliana* atDjBs sequence. **A.** E-values and NCBI accession number of *A. thaliana* atDjBs. **B.** Peptide-binding clefts for Sis1 orthologs. atDjB1, atDjB2, atDjB3, atDjB4, atDjB5, atDjB6, atDjB10 and atDjB17 have peptide-binding clefts similar to Sis1. Surface-filled models of C-terminal region of Sis1 (PDB:1C3G), Sis1 (residues 180 to 352) was made using SWISS MODELER homology-modelling server. Models of peptide-binding clefts of atDjB1 (residues 159 to 335), atDjB2 (residues 173 to 348), atDjB3 (residues 161 to 337), atDjB4 (residues 143 to 323), atDjB5 (residues 172 to 349), atDjB6 (residues 153 to 331), atDjB10 (residues 170 to 347) and atDjB17 (residues 176 to 350) J proteins, based on the crystal structure of Sis1.**C.** Sequence alignment represents the conserved amino acid residues forming the peptide-binding cleft. Conserved amino acids highlighted with colours.

**Fig. S3.**
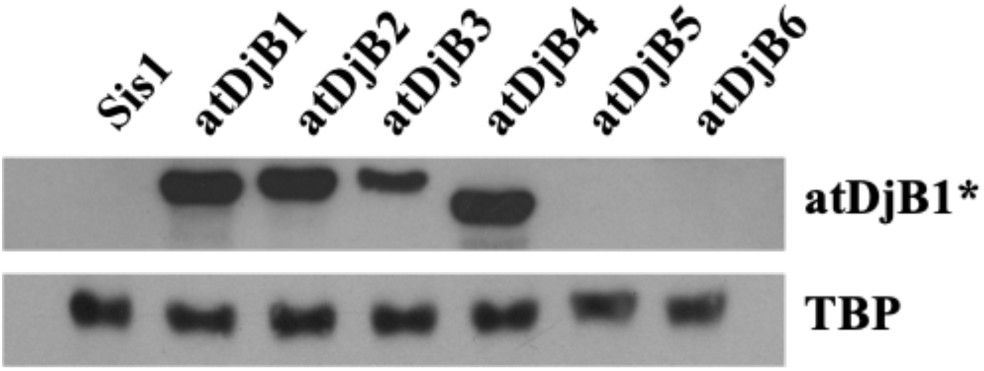
Characterization of a polyclonal antibody against atDjBs. Equal amounts of total cell lysate prepared from s*is1Δ* cells harboring plasmids expressing Sis1 or atDjB1-6 were resolved on SDS-PAGE, electroblotted, and probed with anti-atDjB1 antibody. Anti-TBP1 antibody was used as loading control. Asterisk (*) represent polyclonal antibody against atDjB1 that cross reacts with atDjB2, atDjB3 and atDjB4.

**Fig. S4.**
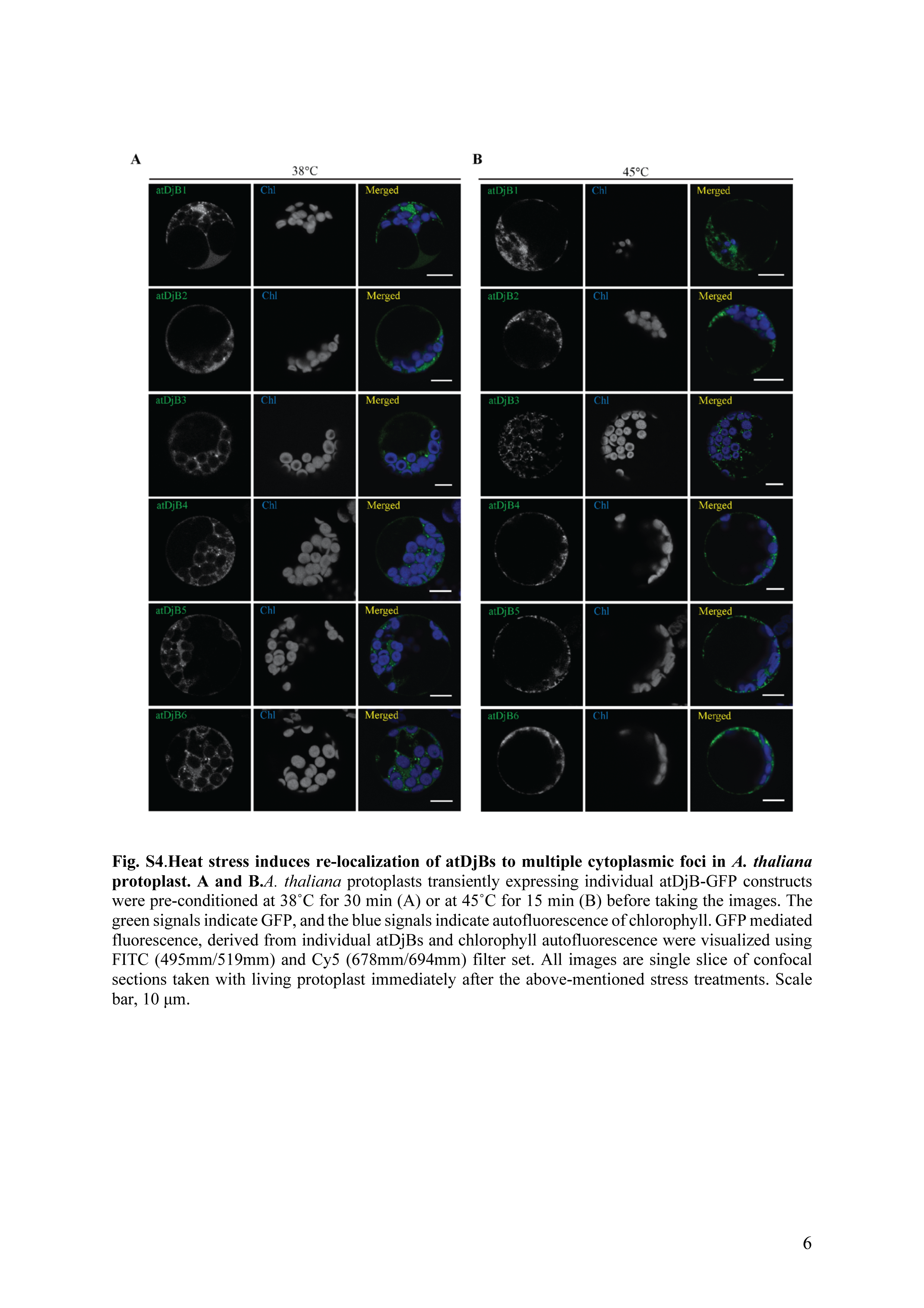
Heat stress induces re-localization of atDjBs to multiple cytoplasmic foci in *A. thaliana* protoplast. **A. and B.** *A. thaliana* protoplasts transiently expressing individual atDjB-GFP constructs were pre-conditioned at 38°C for 30 min (A) or at 45°C for 15 min (B) before taking the images. The green signals indicate GFP, and the blue signals indicate autofluorescence of chlorophyll. GFP mediated fluorescence, derived from individual atDjBs and chlorophyll autofluorescence were visualized using FITC (495mm/519mm) and Cy5 (678mm/694mm) filter set. All images are single slice of confocal sections taken with living protoplast immediately after the above-mentioned stress treatments. Scale bar, 10 μm.

**Fig. S5.**
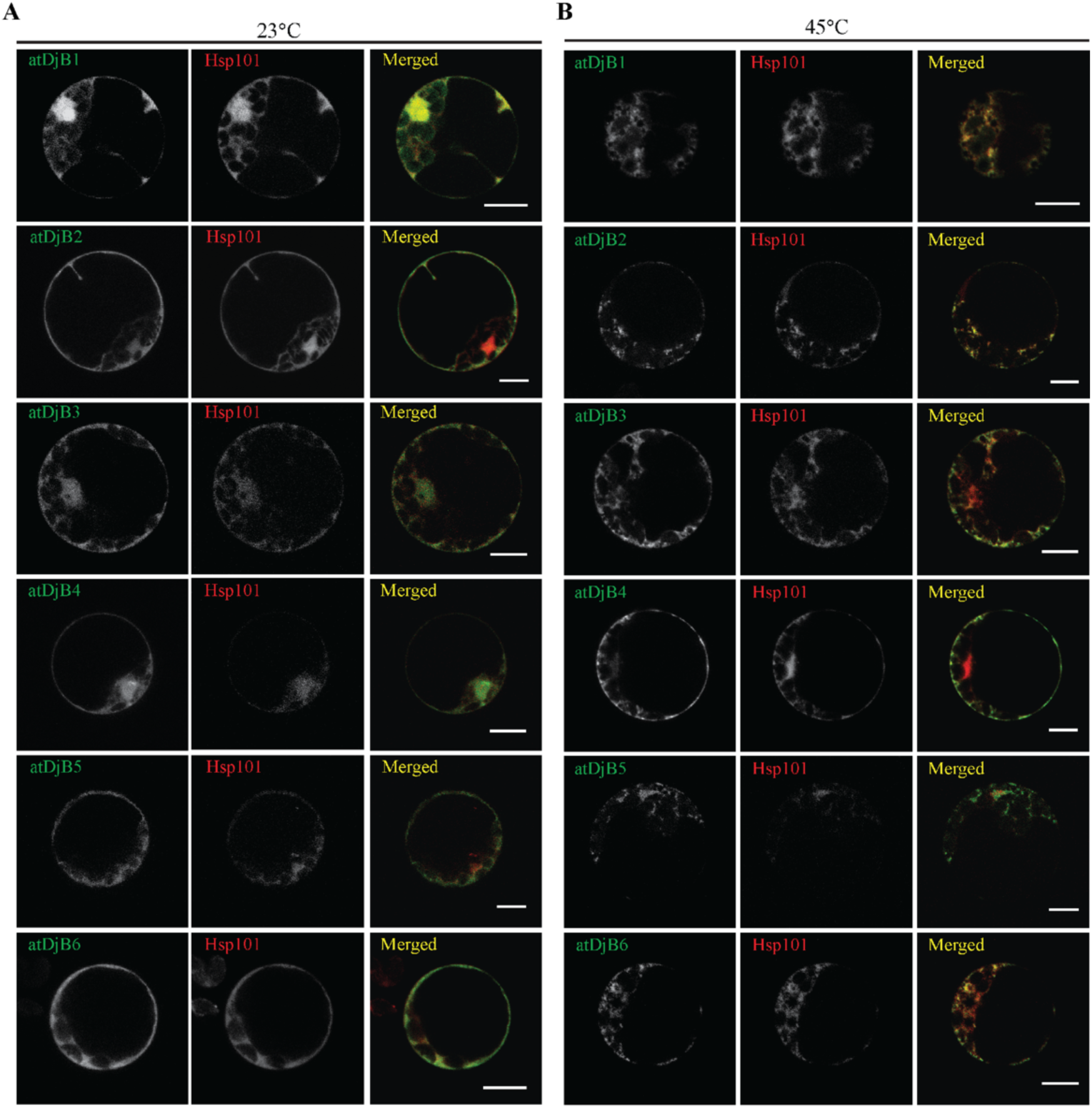
atDjBs colocalize with Hsp101 in *A. thaliana* protoplast. **A. and B.** *A. thaliana* protoplasts transiently expressing individual atDjB-GFP constructs and Hsp101-RFP were visualized at 23°C (A) or after 45°C for 15 min (B) heat stress treatment. Green indicates GFP, red indicates RFP, and blue indicates the autofluorescence of chlorophyll. GFP-mediated fluorescence, derived from individual atDjBs, RFP-mediated fluorescence derived from Hsp101, and chlorophyll autofluorescence were visualized using FITC (495nm/519nm), m-RFP (572nm/606nm) and Cy5 (678nm/694nm) filter set. All images are single slice of confocal sections taken with living protoplast immediately after the above mentioned temperature conditions. Scale bar, 10 μm.

**Fig. S6.**
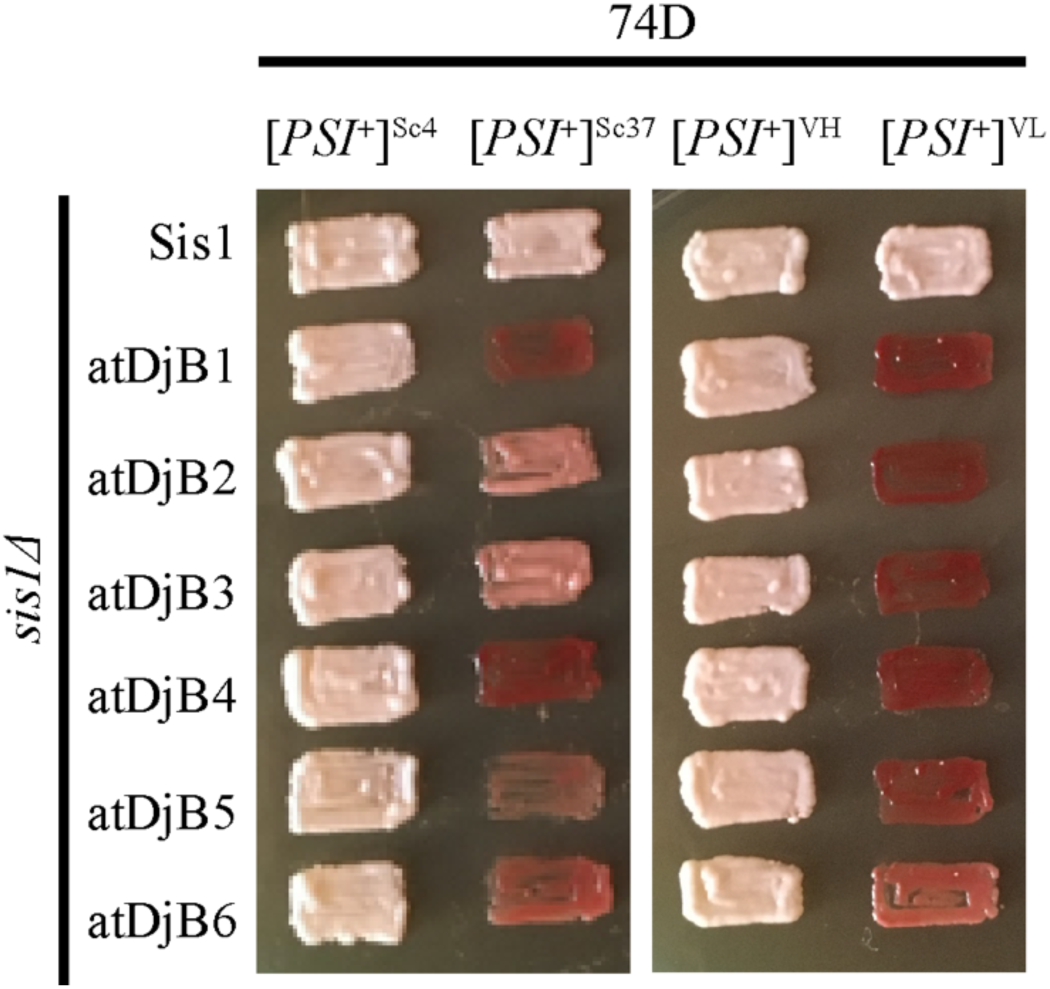
atDjB[*PSI*^+^] maintenance patterns are indistinguishable between the W303 and 74D-694 yeast genetic backgrounds. [*PSI*^+^]^Sc4^, [*PSI*^+^]^Sc37^, [*PSI*^+^]^VH^, and [*PSI*^+^]^VL^*sis1Δ* [*URA3*-*SIS1*] cells were transformed with plasmid (pRS414) expressing Sis1 or atDjB1-6 and subjected to plasmid shuffling on 5-fluorooratic acid (5-FOA). Cells after plasmid shuffling were assayed for [*PSI*^+^] maintenance by colony color on YPD media. Color phenotype assays are shown for representative transformants (n *≥* 10).

**Fig. S7.**
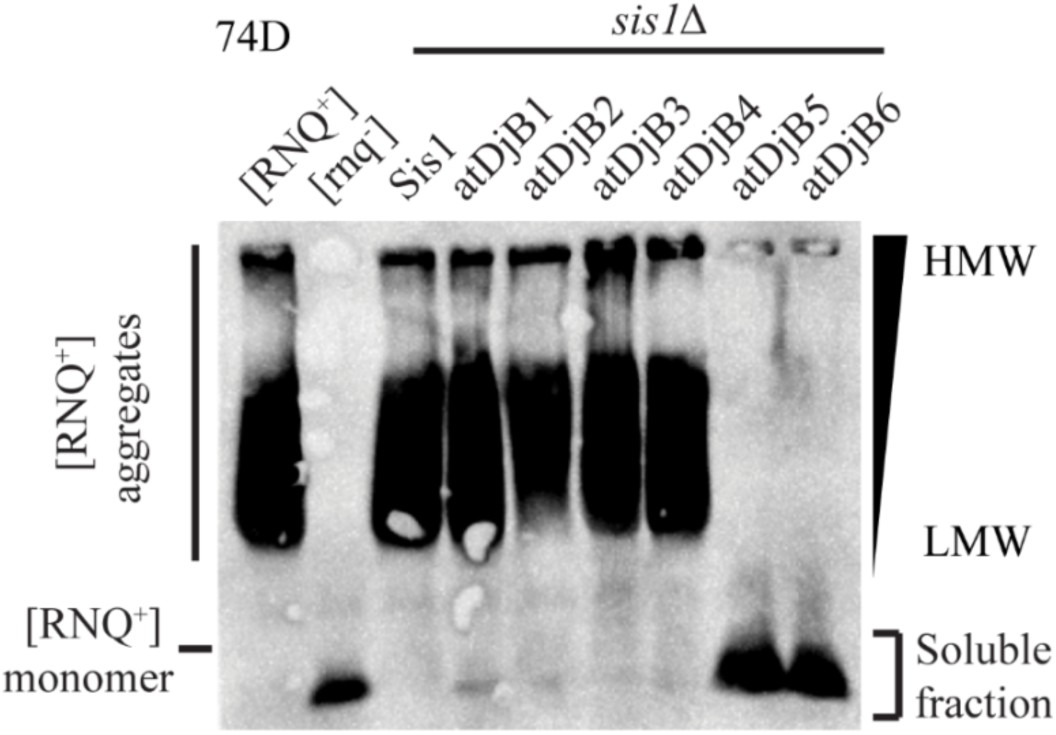
atDjB [*RNQ*^+^] prion maintenance is indistinguishable between W303 and 74D-694 yeast genetic backgrounds. [*RNQ*^+^] *sis1Δ* [*URA3*-*SIS1*] cells of the 74D-694 background were transformed with plasmid (pRS414) expressing Sis1 or atDjB1-6 and subjected to plasmid shuffling on 5-fluorooratic acid (5-FOA). Equal amounts of cell lysate were resolved on semi-denaturing detergent agarose gel electrophoresis (SDDAGE), electroblotted, and probed with anti-Rnq1 antibody. Control [*RNQ*^+^] and GdnHCl-treated [*rnq*^-^] parent cells were included for comparison.

**Fig. S8.**
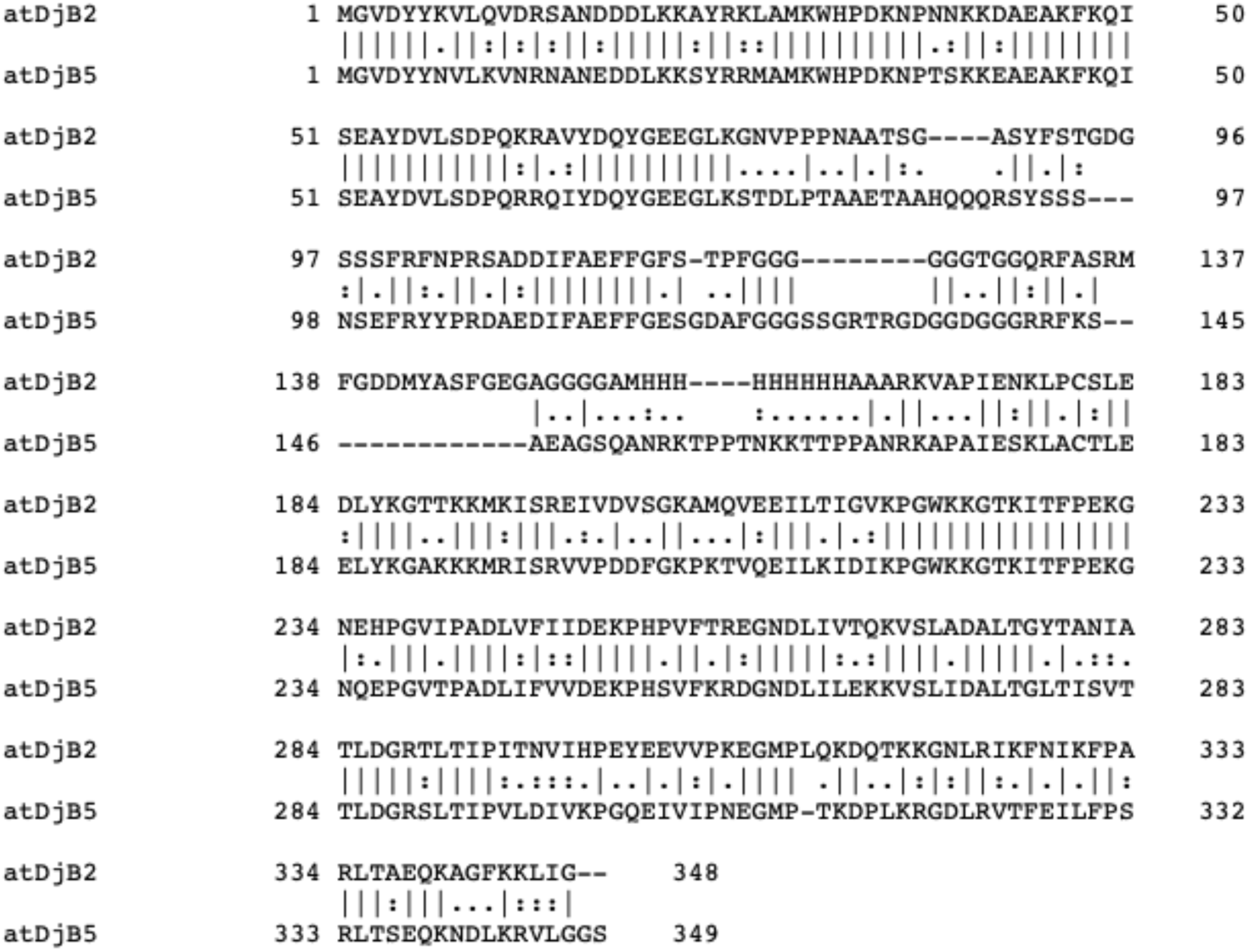
Sequence alignment of the atDjB2 and atDjB5 proteins. Pairwise alignment of amino acid sequences of the atDjB2 and atDjB5 proteins of *A. thaliana* carried out by an online tool EMBOSS Needle that uses the Needleman-Wunsch alignment algorithm.

**Fig. S9.**
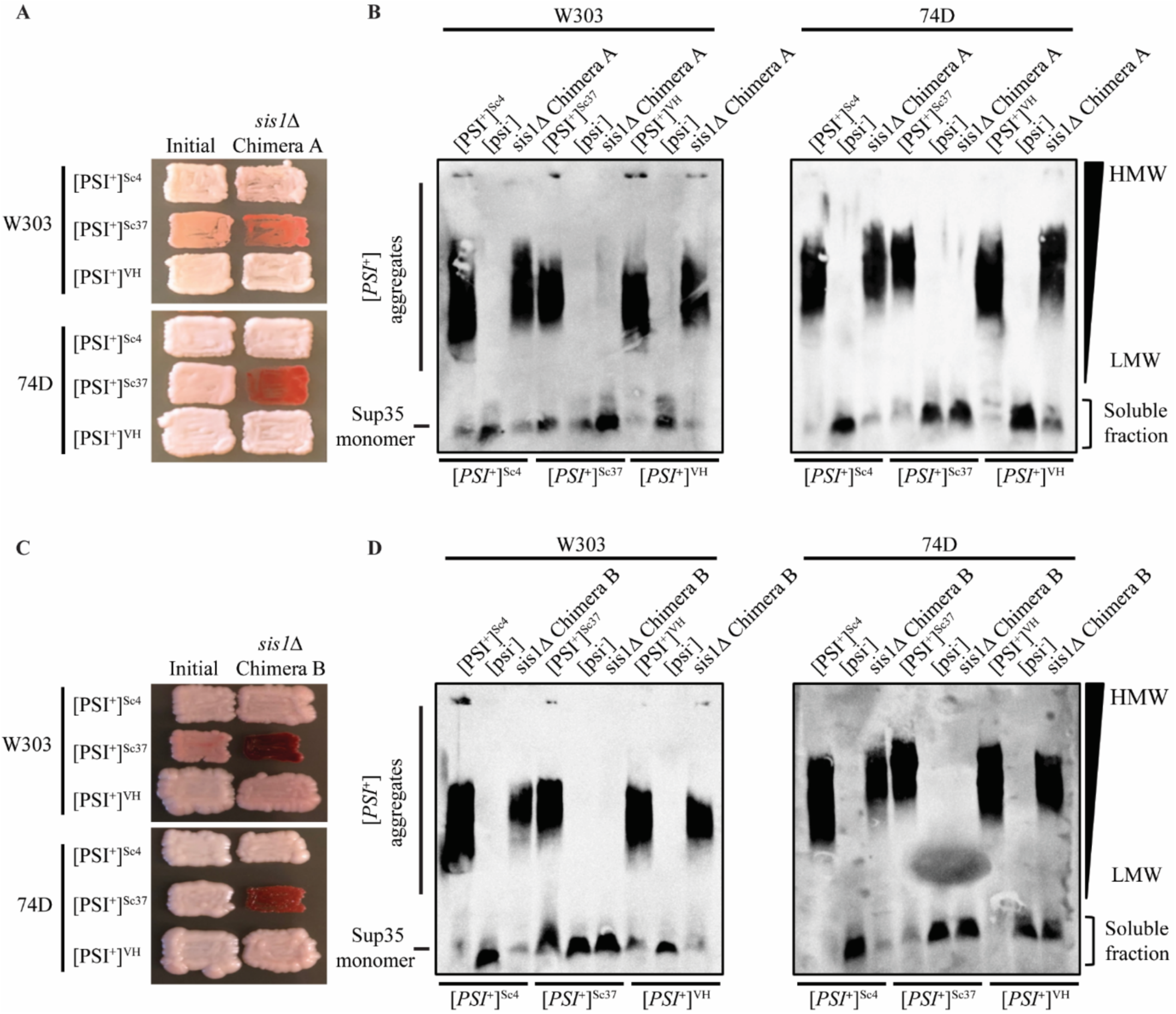
Chimera A and Chimera B [*PSI*^+^] prion maintenance abilities are indistinguishable between W303 and 74D-694 yeast genetic backgrounds. **A and B.** [*PSI*^+^]^Sc4^, [*PSI*^+^]^Sc37^, and [*PSI*^+^]^VH^*sis1Δ* [*URA3*-*SIS1*] cells from two genetic backgrounds, W303 and74D-694, were transformed with plasmid (pRS414) expressing Chimera A and subjected to plasmid shuffling on 5-fluorooratic acid (5-FOA). Cells from individual transformations were assayed for [*PSI*^+^] maintenance by colony color on YPD media (**A**). Equal amounts of cell lysate prepared from shuffled strains were resolved by SDDAGE, electroblotted, and probed with anti-Sup35 antibody (**B**). [*PSI*^+^] and GdnHCl-treated [*psi*^-^] parent cells were included for comparison. **C and D.** [*PSI*^+^]^Sc4^, [*PSI*^+^]^Sc37^, and [*PSI*^+^]^VH^*sis1Δ* [*URA3*-*SIS1*] cells from two genetic backgrounds, W303 and74D-694, were transformed with plasmid (pRS414) expressing Chimera B and subjected to plasmid shuffling on 5-fluorooratic acid (5-FOA). Cells from individual transformations were assayed for [*PSI*^+^] maintenance by colony color on YPD media (**C**). Equal amounts of cell lysate prepared from shuffled strains were resolved by SDDAGE, electroblotted, and probed with anti-Sup35 antibody (**D**). [*PSI*^+^] and GdnHCl-treated [*psi*^-^] parent cells were included for comparison.

**Supporting Table 1.**
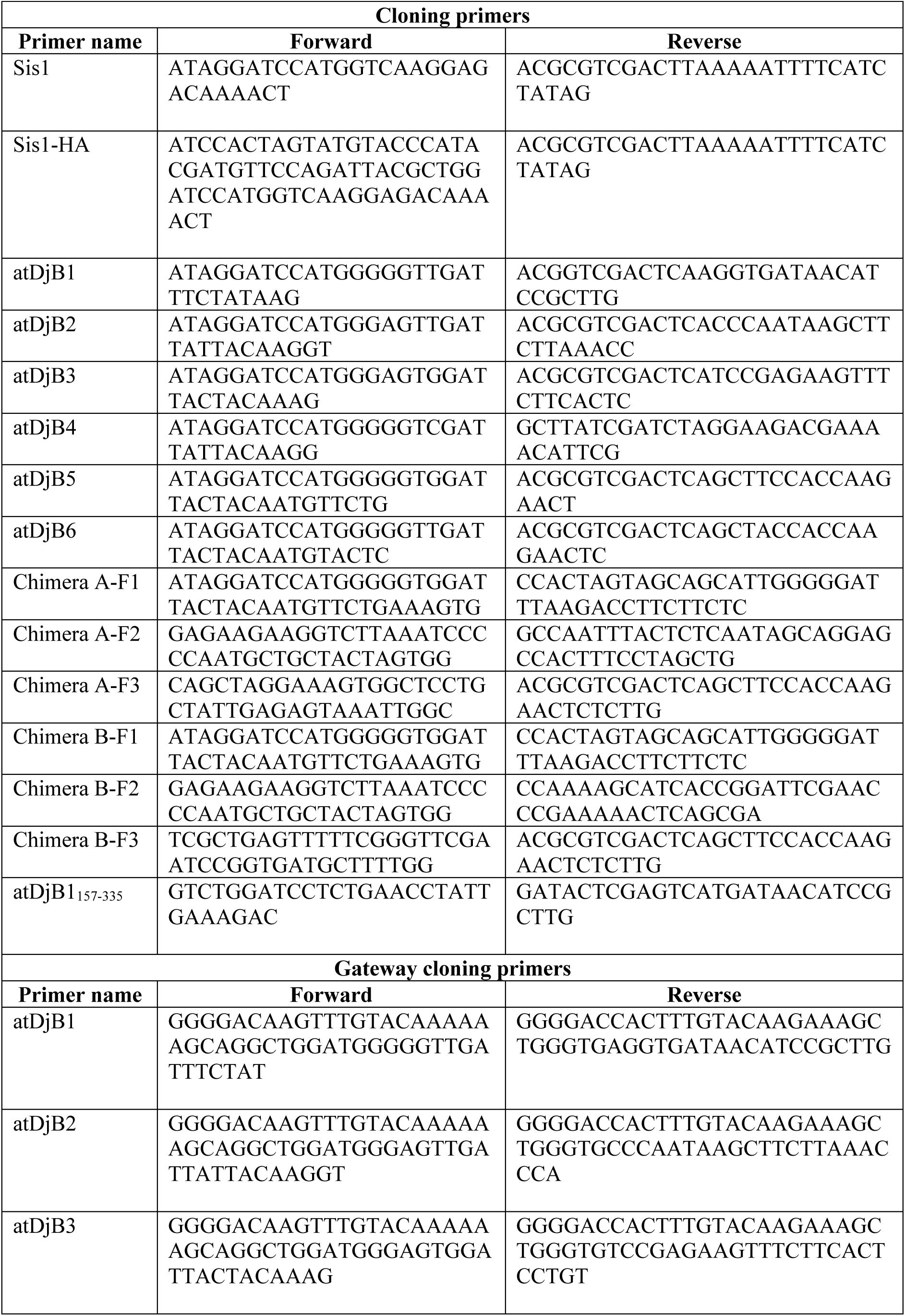
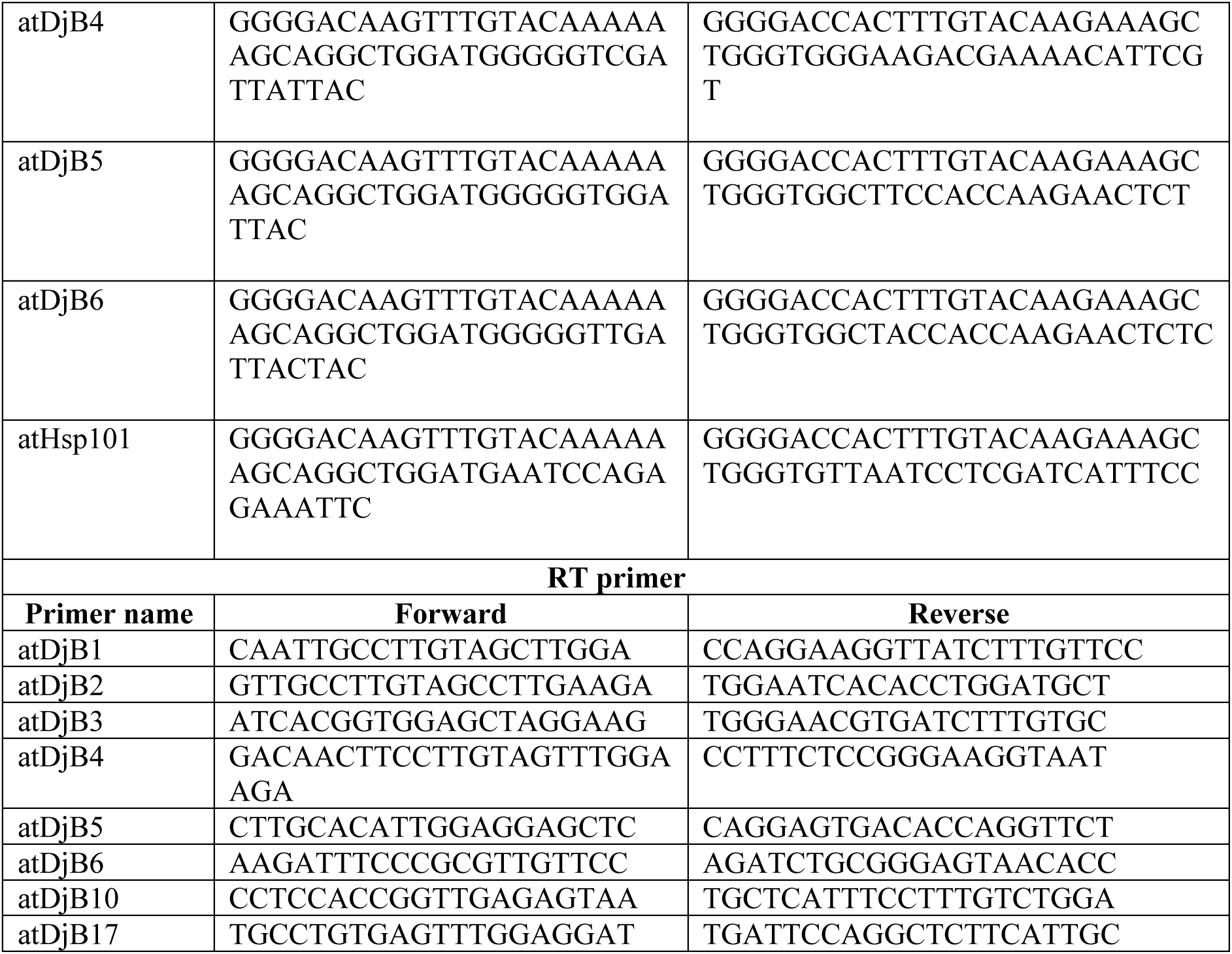
List of Primers used in this work.

